# BLOC: buildable and linkable organ on a chip

**DOI:** 10.1101/2024.10.29.620775

**Authors:** Yusuke Kimura, Hiroki Hamaguchi, Kotaro Oyama, Tomoko G. Oyama, Atsushi Kimura, Yasuhiro Ohshima, Noriko S. Ishioka, Mitsumasa Taguchi

**Author notes:** **Corresponding author:** Mitsumasa Taguchi, National Institutes for Quantum Science and Technology (QST), Takasaki Institute for Advanced Quantum Science., 1233 Watanuki-machi, Takasaki, Gunma, 370-1292, Japan.

## Abstract

Microphysiological systems (MPSs) are attractive alternatives to animal experiments in physiology, pathophysiology, and drug development. However, conventional MPSs are designed for specific applications and have low versatility and scalability. Here, we developed a “Buildable and Linkable Organ on a Chip” (BLOC) to construct diverse MPSs. The Culture, Control, and Analysis BLOCs were standardized to the same size and linked using inlet/outlet joints for perfusion. Radiation-induced chemical reactions were used to suppress the adsorption and absorption of small hydrophobic molecules by BLOCs composed of polydimethylsiloxane. BLOCs for 2D/3D culture could also encapsulate radiation-crosslinked protein hydrogels with designed stiffness and microtopography to mimic the microenvironment *in vivo*. A perfusion device connecting Culture BLOCs was designed for a toxicity assay mimicking the systemic circulation using six types of organoids. An automatic enzyme-linked immunosorbent assay device connecting Culture, Control, and Analysis BLOCs detected proteins released from organoids. BLOCs, which users can freely configure to suit their needs, is expected to become powerful tools in basic research and preclinical trials using MPSs.

## Introduction

Preclinical trials have been conducted in laboratory animals to confirm the safety and effectiveness of pharmaceuticals. However, animal testing is expensive and involves ethical concerns. Additionally, as animal testing does not accurately reproduce human reactions, the failure rate of clinical trials is high, leading to rising medical costs and delays in the development of new effective drugs.^1–3^ Therefore, there is a strong demand for the development of alternative methods to animal testing.

Microphysiological systems (MPSs) have attracted attention as an alternative to animal testing. MPSs are *in vitro* systems that mimic organs or the human body in which cells or organoids are cultured in microfluidic devices that mimic mechanical signals, such as breathing and peristalsis, microenvironments, and blood circulation.^4^ Since Ingber et al. developed a lung-on-a-chip device in 2010,^5^ MPSs that mimic the physiological functions of organs such as the heart, liver, and kidneys, have been reported.^6–8^ Although these devices can evaluate organ-specific physiological functions, they cannot verify the responses of multiple organs to drugs or the effects of organ-to-organ interactions. To solve this problem, MPSs that reproduce multiple organs on a single device have been developed in recent years.^9, 10^ These systems are thought to mimic blood circulation through perfusion and organ-to-organ interactions mediated by metabolites moving in the culture medium. In particular, devices incorporating organoids of various organs derived from human-induced pluripotent stem cells (hiPSCs) are expected to accurately predict pharmacokinetics and toxicity in the human body, which cannot be predicted by animal testing, and are expected to become a new preclinical trial method that will increase the success rate of clinical trials.

However, most of the currently proposed MPSs are fabricated with predesigned functions (e.g., the number, types, and arrangement of culture chambers, connecting channels, and analytical functions), and cannot handle operations outside their design. For example, the strict control over the types of growth and signaling factors in the culture medium required for the generation and culture of organoids^11^ is not possible in existing MPSs with pre-connected culture chambers.^9, 10^ Consequently, the types and numbers of organoids that can be cultured in the existing MPSs are limited. Additionally, the function and fate of cells and organoids are affected by the elastic modulus and surface topography of the culture substrate.^12, 13^ Most conventional MPSs are sealed and cannot encapsulate culture substrates (e.g. hydrogels) suitable to the target cells or organoids. Furthermore, once the design of a conventional MPS is determined, the connection pattern between the culture chambers or the arrangement of the control and analysis systems cannot be changed according to the user’s needs. Consequently, the functions of conventional MPSs are fixed for each device in terms of the cell type, culture method, and analysis purpose, resulting in low versatility and expandability as well as high cost and impracticality.

To develop a practical MPS that solves the above problems, here, we introduce the “Buildable and Linkable Organ on a Chip” (BLOC), which allows users to freely configure functional devices according to their needs. BLOCs are functional chips manufactured to the same size and specifications, with joints at predetermined locations to connect between the BLOCs. Each BLOC has one function, Culture, Control, or Analysis, and users can construct various MPSs by building and linking BLOCs through joints according to their purposes.

Here, we describe the fabrication of the different BLOCs and demonstrate and validate its function. The BLOC is expected to become a powerful tool in basic research and clinical trials utilizing MPSs.

## Results

### Conceptual model of BLOC

To develop a versatile and scalable MPS, we propose BLOC, which allows users to freely configure functional devices according to their requirements. The concept of BLOC is shown in Fig. 1. BLOC was made of polydimethylsiloxane (PDMS),^14, 15^ which is cytocompatible, transparent, and has high oxygen permeability. All the BLOCs were manufactured to the same size and specifications, with joints at predetermined locations to connect the BLOCs and transport fluids between them. Each BLOC was given one function. The functions were classified as a “Culture BLOC” for culturing cells or organoids, a “Control BLOC” for controlling the perfusion and fluid delivery direction of the culture fluid between the BLOCs, and an “Analysis BLOC” for analyzing the components in the culture fluid. Users can build and link BLOCs through joints, for various purposes, including for developing perfusion and cytotoxicity analysis devices. In addition, the Culture BLOC can be coated with the necessary protein reagents or equipped with a suitable culture substrate such as a hydrogel, making it possible to culture various types of cells and organoids in their optimal culture environment. After culturing cells or organoids in an optimal culture environment, a multi-organ MPS can be easily created by connecting the Culture BLOCs. Furthermore, the adsorption and absorption of hydrophobic low-molecular-weight compounds, which are a concern when using PDMS-based BLOCs,^16^ can be reduced without reducing the oxygen permeability or transparency using a modification technique that utilizes radiation-induced reactions.

**Figure 1.**
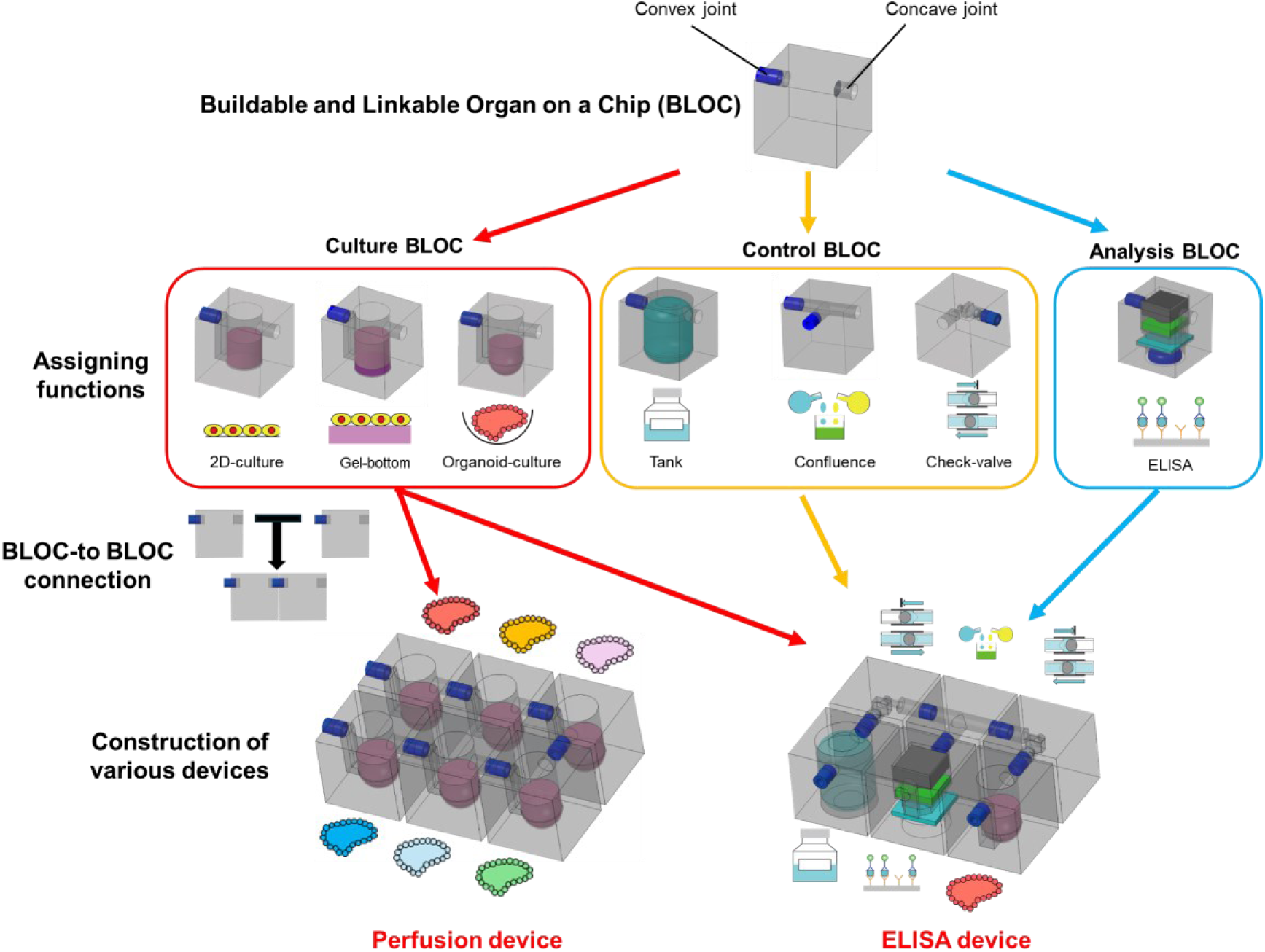
Concept model of the “Buildable and Linkable Organ on a Chip” (BLOC). The BLOCs have standardized size and are linkable by convex and concave joints for perfusion. The BLOCs with various functions are classified as “Culture BLOC,” “Control BLOC,” and “Analysis BLOC”. By connecting the BLOCs, various devices such as perfusion and ELISA devices can be constructed, as demonstrated in this study.

### Basic shape of BLOC

The BLOC was fabricated by assembling multiple PDMS layers using soft lithography techniques. The basic shape of the BLOC is standardized as a block type of 10.0×10.0×16.0 mm (Fig. 2a). The sides of the BLOC are equipped with convex or concave joints. Both types of joints have hollow structures, and liquid transfer between BLOCs is possible by inserting the convex joint of one BLOC into the concave joint of another. In this study, liquid transfer was performed from the convex joint to the concave joint unless otherwise stated. BLOCs are broadly classified into “Culture BLOC,” “Control BLOC,” and “Analysis BLOC” according to their applications, as described below.

**Figure 2.**
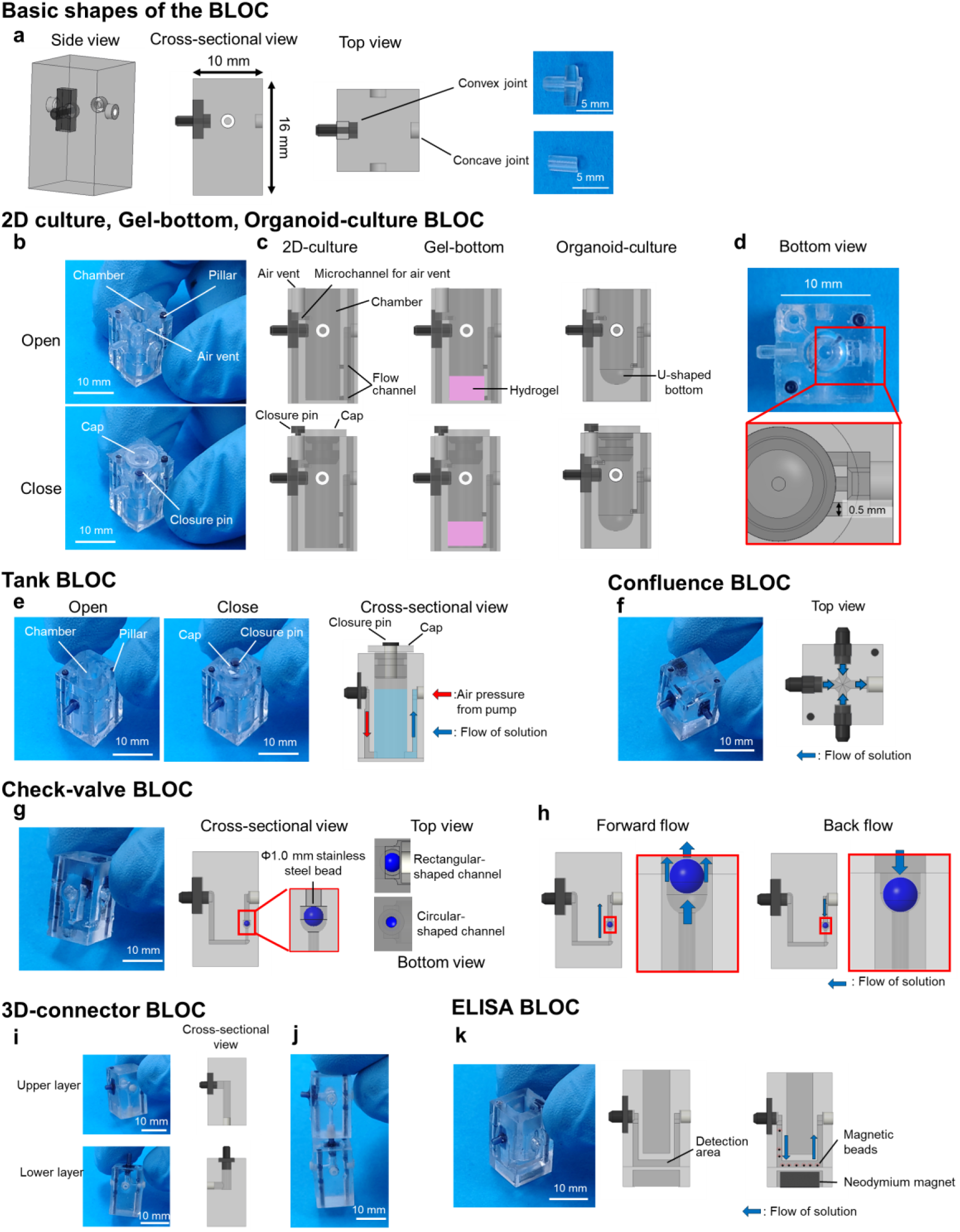
Structure of the BLOC. (a) Basic shapes of the BLOC. (b) Structure of Culture BLOC. Top, Culture BLOC (chamber opened). Bottom, Culture BLOC with the cap and closure pin (chamber closed) (c) Cross-sectional views of the Culture BLOCs. Left, the 2D-Culture BLOC. Center, the Gel-bottom BLOC. Right, the Organoid-culture BLOC. (d) Top, bottom view of the Organoid-culture BLOC. A bottom red frame shows a magnified view of the three microchannels inside the BLOC. (e) Structure of the Tank BLOC. Left, Tank BLOC without (chamber opened) and with (chamber closed) the cap and closure pin. In the chamber of the Tank BLOC, solution is pumped by air pressure through a joint as shown in a right cross-sectional view. (f) Structure of the Confluence BLOC. Left, Confluence BLOC. Right, the top view of the Confluence BLOC. (g) Structure of the Check-valve BLOC. Left, Check-valve BLOC. A right cross-sectional view shows an enlarged view of the Check-valve BLOC and the stainless steel bead in the channel. A rectangular channel (0.7×1.5 mm: top view) was fabricated on the upper side of the bead, and a circular channel (φ0.5 mm: bottom view) on the lower side of the bead. (h) Principle of the Check-valve BLOC. Forward flow is achieved by lifting the beads in a rectangular channel and the solution flows the side of the bead. Back flow pushes the bead toward the circular channel, closing the channel and preventing solution flow. (i) Structure of the 3D-connector BLOC for upper and lower layers. Left, 3D-connector BLOC. Right, a cross-sectional view of the 3D-connector BLOC. (j) Joint Connector BLOCs for upper and lower layers. (k) Structure of the ELISA BLOC. Left, ELISA BLOC. Right cross-sectional views show the microchannel and detection area (3.0×6.0×1.0 mm) of the ELISA BLOC. The ELISA BLOC is equipped with a neodymium magnet to fix magnetic beads only in the detection area.

### Development of Culture BLOCs

In this study, we developed three types of Culture BLOCs: 2D-culture BLOC, Gel-bottom BLOC, and Organoid-culture BLOC (Fig. 2b–d).

#### 2D-culture BLOC

The developed 2D-culture BLOC is shown in Fig. 2b and c. A flat-bottom chamber was fabricated in the BLOC, and it was designed to be able to culture cells in a maximum of 250 µL of culture medium. The BLOC was used for cell culture as is or after plasma treatment to make it hydrophilic. If necessary, the BLOC can also be used after coating with ECM components.

The convex joint of the Culture BLOC was connected to the top of the chamber, and the concave joint was connected to the bottom and center of the chamber via microchannels (Fig. 2b,c). This structure allows solutions to enter the BLOC and flow from the top to the bottom of the chamber, allowing efficient exchange of solutions within the chamber. In this study, a one-inflow, one-outflow-type BLOC equipped with one convex joint and one concave joint was used. However, as shown in Supplementary Fig. 1, BLOCs with different numbers of joints can also be used. By selecting the type of BLOC according to the application, perfusion culture systems with various fluid transfer patterns can be constructed.

The top surface of the BLOC is equipped with a φ1.0 mm air vent that is connected to the top of the chamber through a microchannel. The air pressure generated when the chamber is closed with a cap can be reduced using an open-air vent, preventing leakage of the culture medium and mechanical stress to the cells. The air vent can then be blocked with a closure pin to completely seal off all hole structures, other than the joints.

The microchannel, joint, and air vent structures were also employed in the Gel-bottom BLOC and Organoid-culture BLOC as described below.

#### Gel-bottom BLOC

To culture the cells using ECM-mimicking protein hydrogels, we fabricated a Gel-bottom BLOC (Fig. 2c). It has the same structure as the 2D-culture BLOC described above. Concave joints were connected to the bottom and center of the chamber at two points to prevent the hydrogel formed in the chamber from clogging the flow channel and inhibiting fluid flow. The amount of gel can be adjusted according to the experimental conditions. For example, if 50 µL of gel is formed on the chamber, cells can be cultured in a maximum of 200 µL of culture medium. The gel preparation method is described in detail below.

#### Organoid-culture BLOC

Organoid-culture BLOC has a U-shaped bottom chamber of φ5.0 mm and can culture organoids in up to 200 µL of culture medium (Fig. 2c). The U-shaped bottom positions the organoids in the center of the chamber and prevents them from moving or flowing out, owing to perfusion. The convex joint of the Organoid-culture BLOC was connected to the top of the chamber, and the concave joint was connected to the bottom of the chamber via microchannels. The bottom of the chamber was connected to three Φ0.5 mm microchannels with 0.5 mm spacing; the three microchannels were then integrated into a single microchannel, which was connected to the concave joint (Fig. 2d). This structure prevents ≥0.7 mm size organoids from flowing out to adjacent BLOCs. Arranging the three channels reduces pressure loss and prevents leakage of the delivered solution.

### Development of Control BLOCs

In this study, Tank, Confluence, Check-valve, and 3D-connector BLOCs were developed as follows.

#### Tank BLOC

The developed Tank BLOC is shown in Fig. 2e. The Tank BLOC has a chamber in the center that can hold up to 250 µL of solution. Convex and concave joints were connected to the bottom of the chamber through microchannels. The solution was added to the chamber of the Tank BLOC, which was closed using a cap with an air vent. The chamber was completely closed using a closure pin for the air vents. The solution in the Tank BLOC chamber could be pumped into the connected BLOCs by connecting a convex joint to the exhaust hole side of the ring pump.

#### Confluence BLOC

The developed Confluence BLOC is shown in Fig. 2f. A Confluence BLOC has a simple structure with convex or concave joints mounted on its sides. These joints were connected by a cross-shaped microchannel, which minimized the amount of solution contained in the chips. The solution flowing into one joint was allowed to flow out of the other joints. It was not possible to specify the outflow joints using this chip alone. Therefore, by connecting the Check-valve BLOC described below to each convex joint, we confirmed that the solution flowing in from each convex joint could only flow out to a specific concave joint.

#### Check-valve BLOC

A Check-valve BLOC was developed to precisely control the pumping direction of the solution (Fig. 2g). The Check-valve BLOC has a structure in which the convex and concave joints are connected by a single microchannel. The check-valve structure was mounted on a pathway on the concave side of the joint. A stainless steel bead (φ1.0 mm) was placed in the channel. A circular-shaped channel (φ0.5 mm) was fabricated on the bottom side of the bead, and a rectangular-shaped channel (0.7×1.5 mm) was fabricated on the top side of the bead. When there is no pumping, the bead falls to the bottom due to gravity, and the microchannels are closed. When flow occurs in the desired direction, the microchannel opens with lifting of the bead to the top surface (Fig. 2h). However, when the flow occurs in the opposite direction, the microchannel closes by pushing the bead against the bottom surface.

#### 3D-connector BLOC

3D-connector BLOCs were used to construct multilayered devices. Two types of 3D-connector BLOCs–one for the upper layer and the other for the lower layer–are shown in Fig. 2i. The BLOC for the upper layer has one convex or concave joint mounted on its bottom and side, and that for the lower layer has one convex or concave joint mounted on its top and side. The joints in the BLOC were connected using L-shaped microchannels. As shown in Fig. 2j, the microchannels can be extended not only in the plane but also in the 3D direction by connecting the BLOC for the upper and lower layers via joints.

### Development of Analysis BLOCs

As an example of Analysis BLOCs, we developed an ELISA BLOC (Fig. 2k). The ELISA BLOC has a 3.0×6.0×1.0 mm rectangular detection area at the bottom, with microchannels extending from convex and concave joints on the side of the chip to the detection channel. A 6.0×6.0×2.0 mm neodymium magnet (273 mT) was mounted under the detection area. A solution containing magnetic beads and antibodies flowed into the detection channel through a joint. Because the magnetic beads were fixed to the bottom of the detection area by the magnetic force of the neodymium magnet (Fig. 2k), they did not flow out of the ELISA BLOC, even when the sample and various reaction solutions flowed. The neodymium magnet was removed after adding each solution. The relative amounts of target molecules in the sample were evaluated by analyzing the fluorescence intensity of the beads in the detection channel of the ELISA BLOC under a fluorescence microscope.

### Modification of PDMS utilizing radiation-induced chemical reactions

A concern with the use of the BLOC is that the base material, PDMS, tends to adsorb or absorb small hydrophobic molecules. To overcome this drawback, we attempted to modify PDMS using radiation-induced reactions. We selected highly penetrating γ-rays, because it is capable of modifying the entire bulk of a BLOC measuring 10.0×10.0×16.0 mm as well as modifying many BLOCs at once. Fig. 3 summarizes the evaluation of the chemical and mechanical properties of the PDMS before and after γ-ray irradiation.

**Figure 3.**
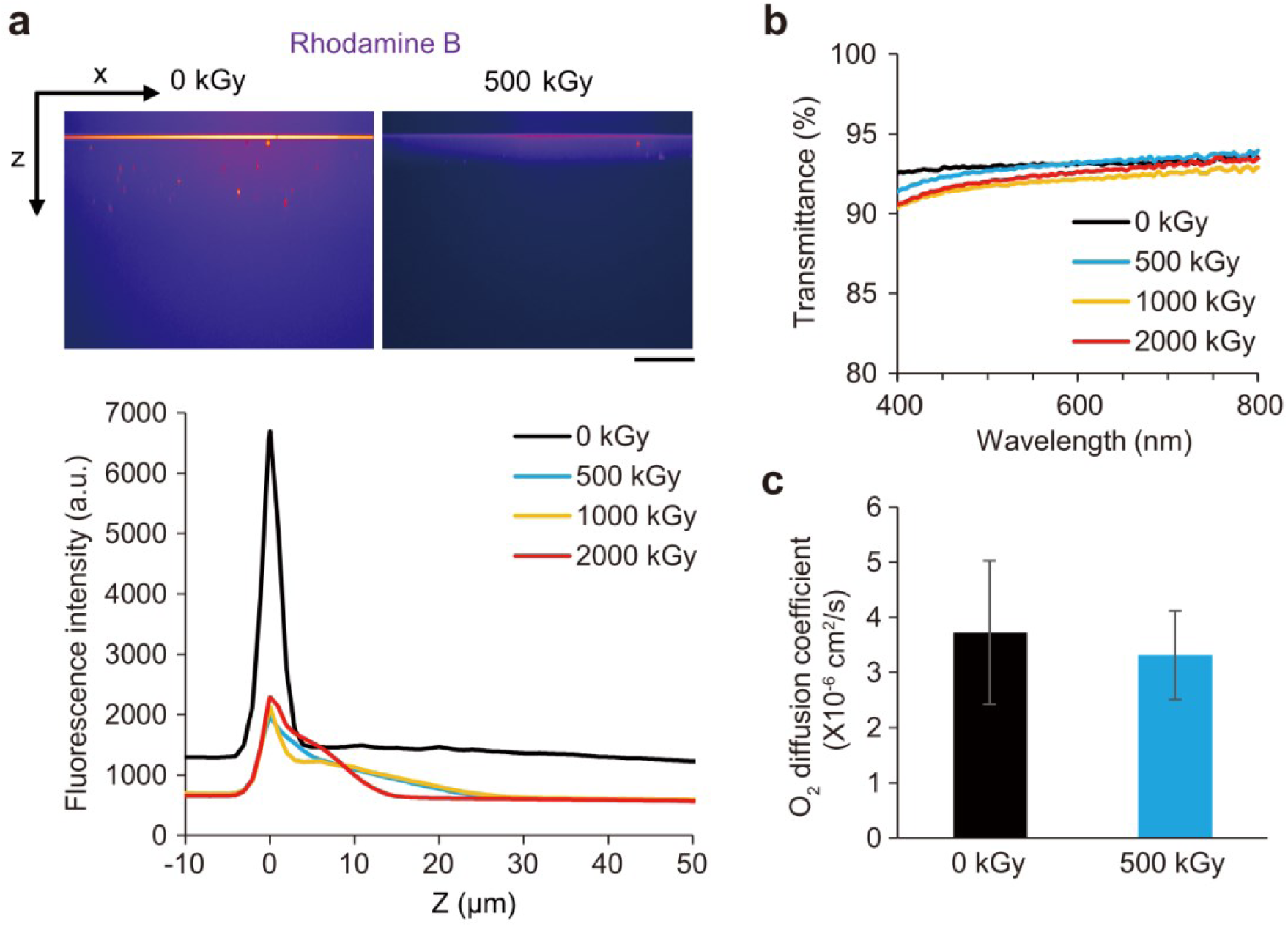
Modification of polydimethylsiloxane (PDMS) utilizing radiation-induced chemical reactions. (a) Top, cross-section images of rhodamine B in the non-irradiated or 500 kGy ^60^Co γ-ray-irradiated PDMS substrates obtained by confocal fluorescence microscopy. Scale bar, 50 μm. Bottom, depth distribution of rhodamine B. (b) Ultraviolet-visible spectra of PDMS substrates before and after irradiation with γ-rays. (c) Diffusion coefficient of O_2_ in PDMS substrates before and after irradiation with γ-rays. *N* = 3. Error bars, SEM. *P*-value from two-sided t-test between 0 kGy vs. 500 kGy is 0.803.

The adsorption and absorption of small hydrophobic molecules were evaluated using rhodamine B as a model drug. The color map of the fluorescence intensity in the cross section of the samples represents the distribution of rhodamine B (Fig. 3a). Significant adsorption was observed near the surface of the non-irradiated PDMS, and the dye attached to the sample surface was observed as a bright horizontal line. In addition, the non-irradiated sheets clearly absorbed rhodamine B. In contrast, both the adsorption and absorption of rhodamine B were remarkably suppressed after the γ-ray irradiation. Irradiation with 500 kGy clearly suppressed the adsorption/absorption of rhodamine B, and the inhibitory effect was not significantly affected by irradiation doses up to 2,000 kGy.

Next, the effect of γ-ray irradiation on the transparency of PDMS was evaluated at wavelengths of 400–800 nm, which are used for observing cells (Fig. 3b). Although the transparency decreased slightly with increasing irradiation dose, it remained comparable to the transparency of conventional polystyrene cell culture dishes (>90%) within a dose range of 500–2,000 kGy.

Considering the suppression of adsorption/absorption properties and maintenance of transparency, it was determined that an irradiation dose of 500 kGy was optimal for modifying the BLOCs in this study. We investigated the effect of this modification method on the oxygen permeability, which is particularly necessary for culturing organoids and cell aggregates. Gas permeation curves were obtained by plotting the oxygen concentration permeating the PDMS sheets over time. The oxygen diffusion coefficient calculated from the gas permeability curves showed no significant differences before and after modification (Fig. 3c), indicating that the modified PDMS retained high oxygen permeability. Furthermore, modification by γ-ray irradiation does not require chemicals. Thus, this modification method suppresses the adsorption/absorption of the hydrophobic small molecules of BLOC while retaining the advantages of PDMS, such as transparency, high oxygen permeability, and biocompatibility.

### BLOCs for cell culture mimicking *in vivo* physiological conditions

We investigated whether the BLOC prepared and modified using the above method was suitable for cell culture. Human skin squamous cell carcinoma (HSC-1) cells adhered to the BLOC with plasma treatment and survived for at least 48 h (Supplementary Fig. 2), confirming that the BLOC enables conventional 2D culture.

However, the measured elastic modulus of PDMS, the base material of the BLOC, was approximately 5 MPa, which is much higher than the reported range of the elastic modulus of soft tissues *in vivo* (from ∼1 to a few hundreds of kPa).^17^ Cells are affected not only by their chemical composition but also by the elasticity and topography of the ECM.^18–20^ To reduce *in vivo* and *in vitro* behavioral disparity of cells,^21–23^ it is necessary to encapsulate a protein hydrogel that can provide an environment closer to the *in vivo* ECM within the Culture BLOC. We attempted to produce a Gel-bottom BLOC that chemically and mechanically mimics the *in vivo* ECM. To create the hydrogel, we applied our technology to crosslink proteins without using chemical crosslinkers and controlled the stiffness and microstructure of the gel.^24^

The hydrogel was made from gelatin, a hydrolysate of type I collagen, which is the main component of *in vivo* ECM. A gelatin solution was injected into the Gel-bottom BLOC, cooled to form a physical gel, and irradiated with γ-rays. Irradiation with a dose of 8–25 kGy altered the physical hydrogel into crosslinked hydrogels without any chemical treatment (Fig. 4a). The crosslinked gelatin hydrogels are insoluble at 37 °C, so they can be used for cell culture. The elastic moduli of the hydrogels were regulated by adjusting the irradiation dose (Fig. 4b). A higher irradiation dose resulted in higher crosslinking density and elastic modulus. The obtained hydrogel covered a broad range of *in vivo* elasticities of soft tissues.

**Figure 4.**
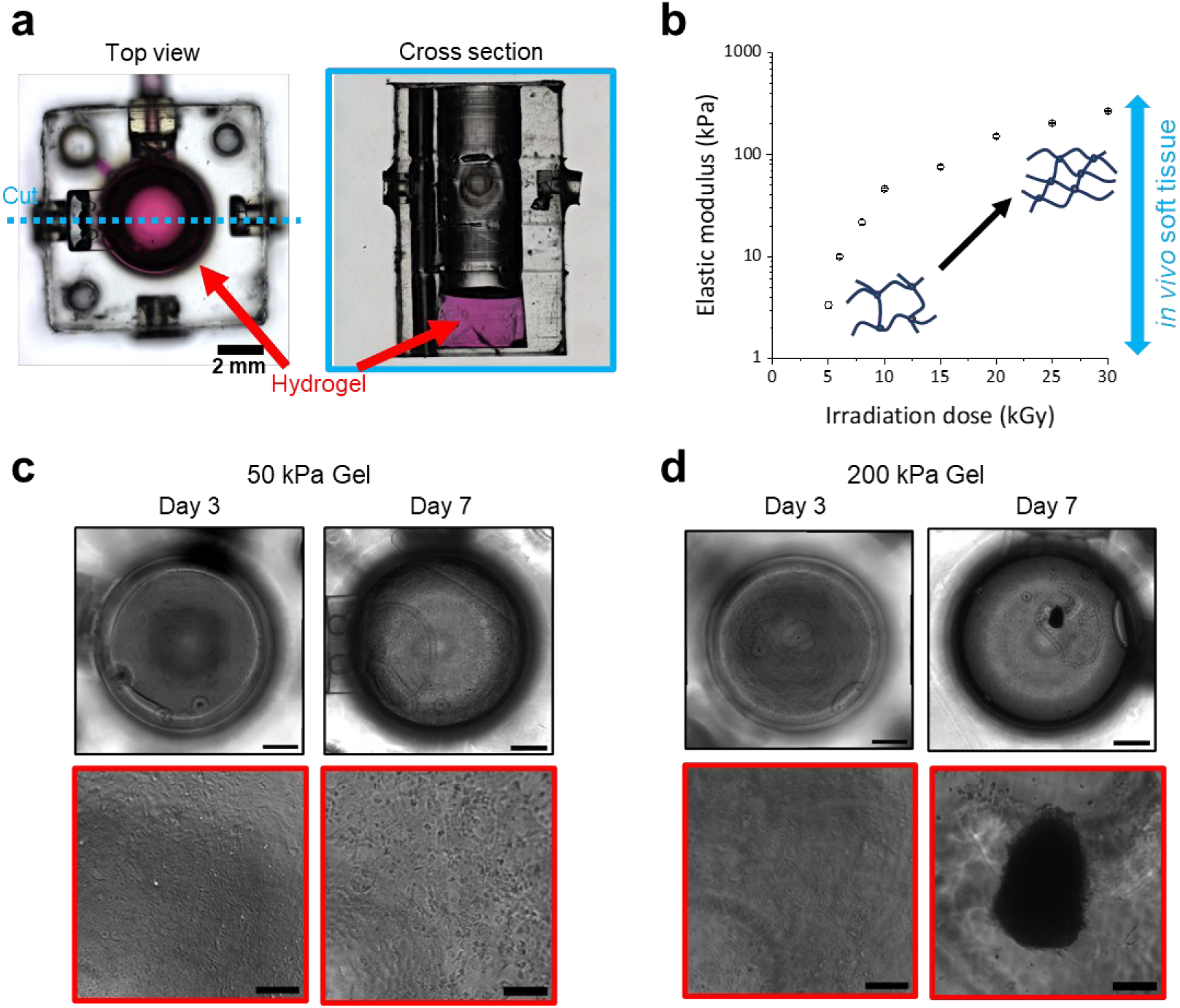
Extracellular matrix (ECM)-mimicking gelatin hydrogel encapsulated in the Gel-bottom BLOC. (a) Left, bright-field image taken from the top of the ECM-mimicking gelatin hydrogel fabricated in the Gel-bottom BLOC. Right, bright-field image taken from the cross section of the hydrogel in the BLOC. (b) Elastic modulus of the gelatin hydrogel. The crosslinking density of gelatin molecules increases with the irradiation dose of γ-rays, resulting in stiffening of the hydrogel. The double arrow line indicates the range of elastic modulus in soft tissue *in vivo* (from ∼1 to a few hundreds of kPa). *N* = 3. Error bars, SEM. (c, d) HSC-1 cells cultured for 7 days in the Gel-bottom BLOC containing 50 kPa (c) or 200 kPa (d) hydrogel. Left, phase contrast images on the third day. Right, phase contrast images on the 7^th^ day. The bottom images show the enlarged view of the region marked in red squares in top images. Scale bars in the top and bottom images are 1000 μm and 200 μm, respectively.

The cells were cultured in the fabricated Gel-bottom BLOC. In the case of HSC-1 cells, a monolayer morphology was observed on hydrogels with a stiffness of 50 kPa, regardless of the culture period (Fig. 4c). In contrast, HSC-1 cells formed spheroids on the 230 kPa hydrogel, depending on the culture period (Fig. 4d). This indicates that HSC-1 cells respond to a certain stiffness and adopt a morphology similar to that of tumors *in vivo*. This further suggests that the cell morphology can be controlled by the stiffness of the hydrogel.

Additionally, the hydrogel surface can be processed to obtain microstructures as soft as *in vivo* ECM. Supplementary Fig. 3a shows microgrooves with widths and spacing of 50 µm and a height of 2 µm, formed on the hydrogel surface. C2C12 myotubes cultured on the hydrogel with microgrooves were aligned similarly to the skeletal muscle *in vivo*, unlike those cultured on a flat hydrogel (Supplementary Fig. 3b).

Thus, by incorporating ECM-mimicking hydrogels with controlled elasticity and microtopography into BLOCs, cells can be induced to adopt a morphology similar to that *in vivo*.

### Device construction and fluid delivery test

By connecting BLOCs that were fabricated and modified using the methods described above, various devices tailored for the intended use were constructed. A bottom frame was prepared in advance, in which 10 mm square lattice structures with a φ7.5 mm hole in the center were arranged at 1.5 mm intervals. Various devices were constructed by installing a BLOC in the bottom frame (Fig. 5a,b). This installation method allowed the position and spacing of each BLOC to be fixed accurately, preventing leakage between the joints during liquid transfer. After connecting all the BLOCs, caps and closure pins were attached to the BLOCs as necessary (Fig. 5c). Furthermore, BLOCs were firmly fixed in place by attaching an upper frame made of 10 mm square lattice structures with a φ5.5 mm hole in the center. This upper frame allows the BLOC to be completely fixed in position and prevents its disassembly during device operation. Furthermore, this could hold down the caps of the BLOCs, preventing leakage of the solution during long-term perfusion culture (Fig. 5d,e).

**Figure 5.**
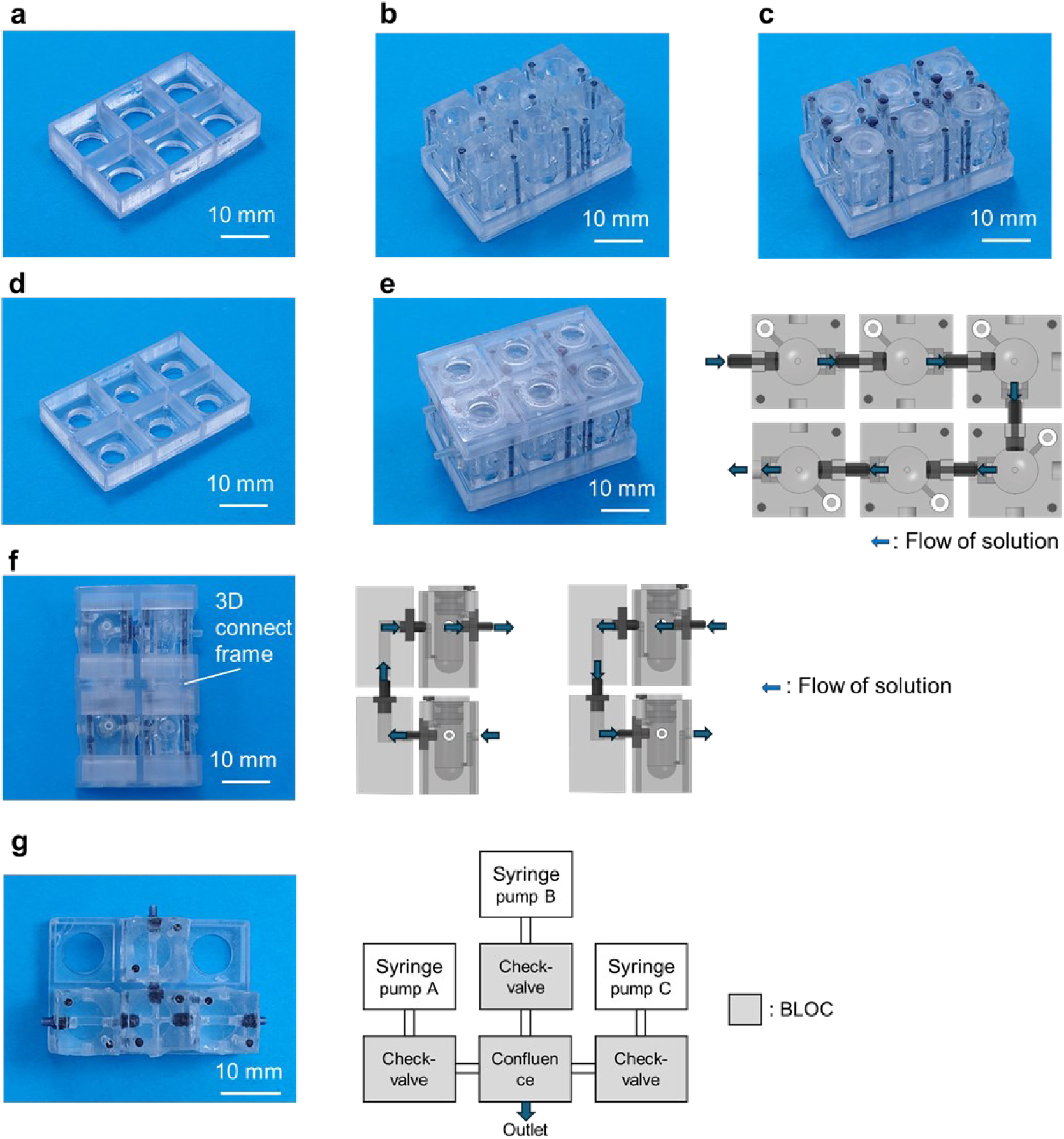
Construction of BLOC device. (a) Bottom frame of the 2D-perfusion device. (b) Constructed 2D-perfusion device without cap and closure pin (chamber opened). (c) Constructed 2D- perfusion device with cap and closure pin (chamber closed). (d) Upper frame of the 2D-perfusion device. (e) Constructed 2D-perfusion device with upper frame. The right figure shows the flow of liquid supply. (f) Constructed 3D-perfusion device. The right figures show the flow of liquid supply. (g) Device for verifying the Check-valve BLOC operation. Left, the picture of the constructed check-valve device. Right, the structure of check-valve device and control system.

We tested whether the devices constructed by connecting the BLOCs could transport liquid as designed by using two types of devices: a 2D perfusion device in which Culture BLOCs were connected in a planner manner (Fig. 5e), and a 3D perfusion device in which Culture BLOCs were connected three-dimensionally with 3D connector BLOCs (Fig. 5f). We tested the delivery of colored water at 100 µL/min using a syringe pump connected to the inlet via a silicone tube. Colored water was delivered to all BLOCs in the device without any leakage from the joints or caps (Supplementary Movie 1). Next, one Culture BLOC filled with colored water was removed from the 2D perfusion device and connected to a different 2D liquid transport device. By applying a positive pressure of 100 µL/min using a syringe pump, the colored water in the BLOC could be delivered to the other device (Supplementary Movie 2). In the case of the 3D perfusion device, it was confirmed that colored water could be transferred from the lower to the upper BLOCs and vice versa (Supplementary Movie 3).

We constructed a device to verify the operation of the Check-valve BLOC (Fig. 5g). When three colored waters were pumped at 100 µL/min through three Check-valve BLOCs to one Confluence BLOC, each colored water was pumped in only the intended direction (Supplementary Movie 4). These results demonstrate that, by connecting the developed BLOCs, it is possible to construct devices that can precisely transport liquids in the desired direction in two and three dimensions.

### Toxicity assay of human organoids in the perfusion device mimicking systemic circulation

We designed a toxicity assay using a BLOC-based perfusion device for six types of hiPSC-derived organoids (stomach, intestine, liver, heart, brain, and kidney), mimicking oral intake and systemic circulation (Fig. 6a). As shown in Fig. 6a, six Organoid-culture BLOCs were connected, and three organoids of one type were placed in each BLOC. Perfusion medium containing toxic agent (30 mM acrylamide) and two fluorescent dyes, Hoechst for total cells and propidium iodide (PI) for dead cells, was flowed at 10 µL/min using a syringe pump from the stomach to the kidney for 24 h, and the fluorescence images were captured by a CMOS camera every hour (Fig. 6b,c; Supplementary Movies 5 and 6). During perfusion for 24 h without a toxic agent, the ratio of the intensity of PI to that of Hoechst was relatively stable (Fig. 6c,d). However, in the presence of toxic agents, the PI/Hoechst values of the liver, heart, brain, and kidney were increased significantly compared to those of the control (Fig. 6b–d). These results demonstrate that this BLOC-based perfusion device is functional for both co-culture and toxicity assays of multiple organoids under conditions that mimic systemic circulation.

**Figure 6.**
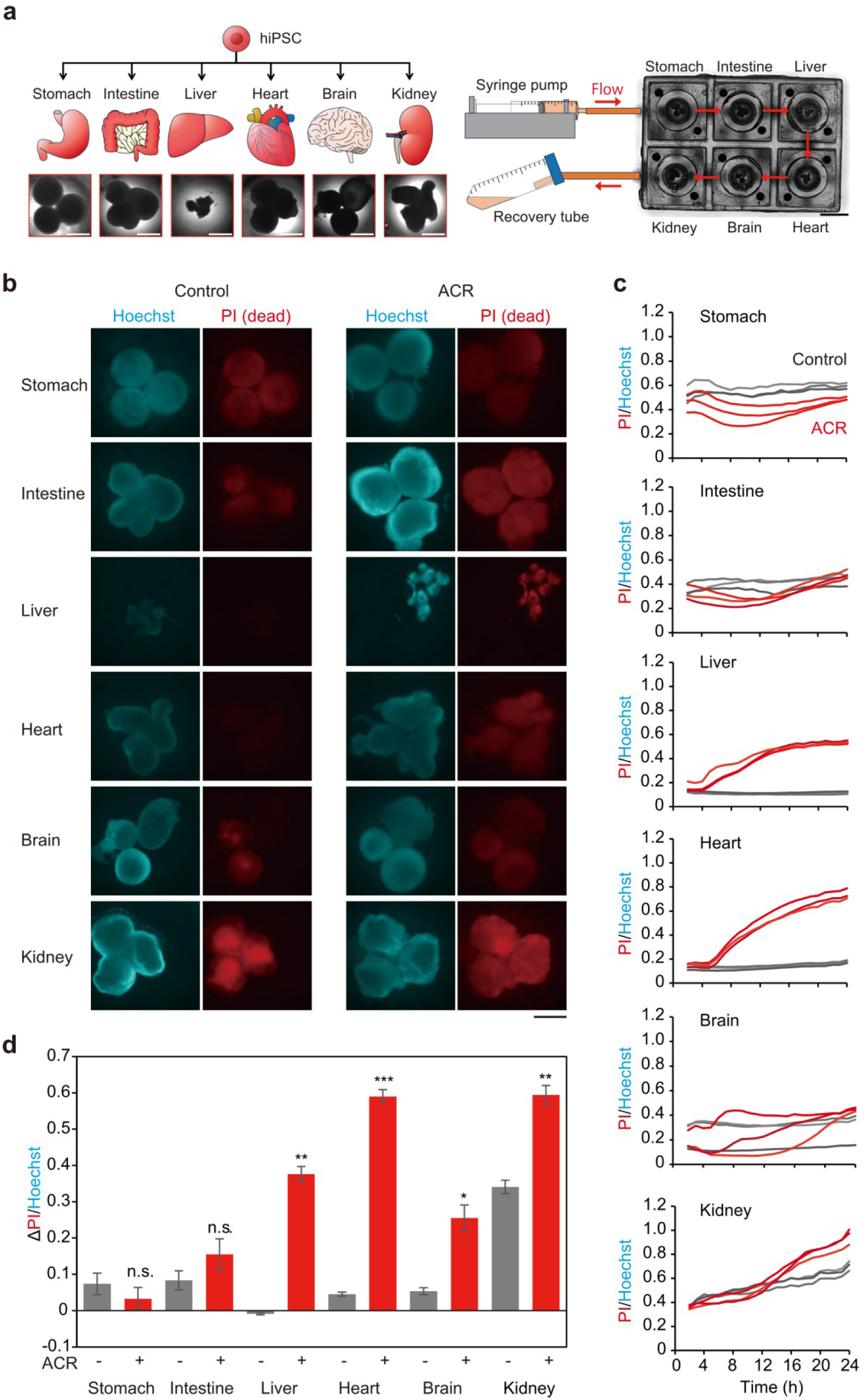
Toxicity assay of human organoids connected in the perfusion device. (a) Left, bright field images of six kinds of organoids differentiated from human induced pluripotent stem cells (hiPSCs) used in the toxicity assay. Right, schematic illustration of toxicity assay of organoids connected in the perfusion device. Organoid-culture BLOCs for different organ organoids are connected by inlet/outlet joints. Culture medium containing fluorescent dyes Hoechst for total cells and propidium iodide (PI) for dead cells is flowed at 10 µL/min by syringe pumps from stomach to kidney as red arrows indicated. Fluorescence images were captured by a CMOS camera every hour. White scale bars, 1 mm. Black scale bar, 5 mm. (b) Fluorescence images of six kinds of organoids stained by Hoechst (cyan) and PI (red) at 24 h. Left, control without a toxic agent; Right, 30 mM acrylamide (ACR). Days of differentiation are 48, 48, 21, 19, 78, and 25 for the stomach, intestine, liver, heart, brain, and kidney organoids, respectively. Scale bar, 1 mm. (c) Ratio values of fluorescence intensities of PI divided by Hoechst (PI/Hoechst) of each organoid at different time points. Black and red lines are PI/Hoechst for control and 30 mM ACR, respectively. (d) The changes in PI/Hoechst from 2 h to 24 h (ΔPI/Hoechst) for each organoid. Black and red boxes indicate control and 30 mM ACR, respectively. Error bars, SEM. *P*-values from two-sided t-test between control vs. ACR are 0.48, 0.32, 0.0038, 7.6×10^-4^, 0.038, and 0.0041 for stomach, intestine, liver, heart, brain, and kidney, respectively. *, *P*<0.05; **, *P*<0.01; ***, *P*<0.001; n.s., not significant (*P*>0.05).

### Automated ELISA device detects proteins released from organoids

Finally, we constructed an ELISA device by connecting the Culture, Control, and Analysis BLOCs. To functionally evaluate the ELISA device, we performed a protein detection assay on the organoids. Cells release non-secreted proteins when damaged by drugs. Therefore, we investigated whether the toxicity of acrylamide to heart organoids could be evaluated based on the relative amount of GAPDH released into the culture supernatant upon cell damage^25^ using the constructed ELISA device. Heart organoids were placed in Organoid-culture BLOC and cultured in control medium or acrylamide-containing medium at 37 °C for 24 h. After incubation, three Tank BLOCs, three Check-valve BLOCs, one Confluence BLOC, and one ELISA BLOC were connected to construct the ELISA device (Fig. 7a). Three ring pumps were connected to the Tank BLOC and the Organoid-culture BLOC, and the pumping speed and timing of each ring pump were controlled by a program to automate the antigen-antibody reaction and washing procedure in the ELISA device.

**Figure 7.**
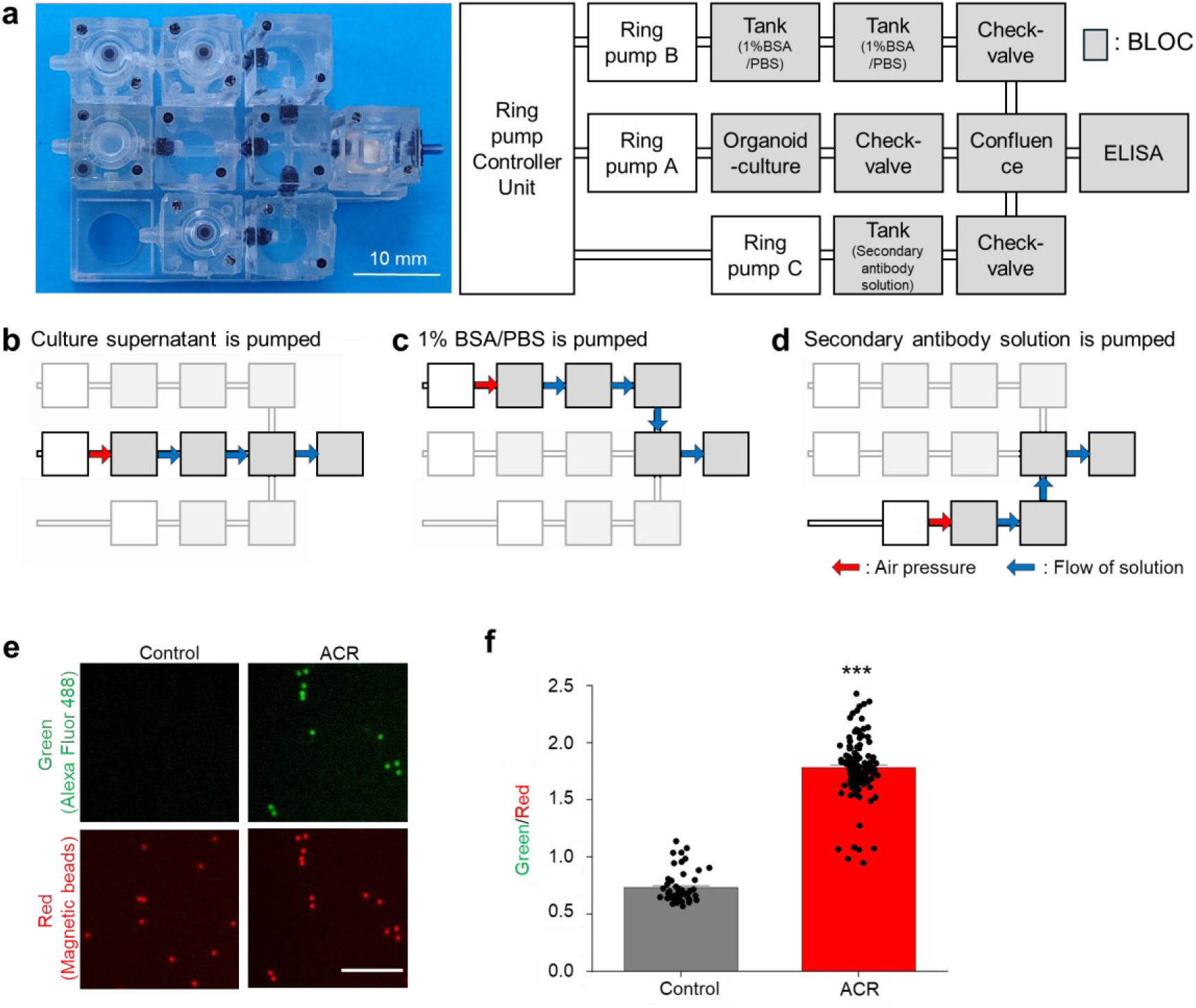
Automated enzyme-linked immunosorbent assay (ELISA) device for detecting protein released by organoids. (a) Left, the picture of the constructed ELISA device. Right, the structure of ELISA device and control system. (b)–(d) How to use the ELISA device. (b) The culture supernatant in the Organoid-culture BLOC is pumped to the ELISA BLOC to allow binding of the target protein to the primary antibody-conjugated red-fluorescent magnetic beads. (c) 1% BSA in PBS is pumped to the ELISA BLOC for ejecting culture supernatant or secondary antibody solution. (d) 1% BSA in PBS containing green-fluorescent secondary antibody is pumped into the ELISA BLOC to allow the binding of the secondary antibody to the target protein captured on the magnetic beads in the ELISA BLOC. (e) Fluorescence images of Alexa Fluor 488 conjugated with GAPDH antibody (green) and magnetic beads conjugated with GAPDH antibody (red) in ELISA BLOC after the reaction with supernatant of heart organoids cultured in the Organoid-culture BLOC for 24 h without toxic agent (Control in left) or with 30 mM acrylamide (ACR in right). Scale bar, 100 µm. (f) The ratio intensity of Alexa Fluor 488 (green) to magnetic beads (red). *N* = 52 and 126 beads for control and with 30 mM ACR, respectively. Error bars, SEM. *P*-value from two-sided t-test between control vs. ACR is 3.3×10^-78^. ***, *P*<0.001.

We conducted ELISA using the following sequence: the culture supernatant was supplied from the Organoid-culture BLOC to the ELISA BLOC and left to stand at room temperature (∼25 °C) for 2 h (Fig. 7b). This allowed the GAPDH present in the supernatant to bind to the primary antibody-conjugated magnetic beads in the ELISA BLOC. The ELISA BLOC was washed with 1% BSA in PBS supplied by the Tank BLOC (Fig. 7c). Subsequently, the secondary antibody solution was supplied by the Tank BLOC to the ELISA BLOC and left to stand at room temperature for 1 h (Fig. 7d). During this process, secondary antibodies were bound to the GAPDH captured by the beads. Finally, the ELISA BLOC was washed with 1% BSA in PBS supplied by the Tank BLOC (Fig. 7c). The ELISA device performed complex pumping operations using multiple types of solutions. Each solution could be pumped accurately without backflow or contamination owing to the Check-valve BLOC.

After the above sequence, the ELISA BLOC was removed from the device, and the red fluorescence of the magnetic beads and the green fluorescence of the secondary antibodies were photographed under a fluorescence microscope (Fig. 7e). From the obtained images, the ratio of red to green fluorescence, that is, the relative amount of GAPDH, was calculated (Fig. 7f). A significant increase in the relative amount of GAPDH was observed in the supernatants of heart organoids cultured with acrylamide. These results indicated that the BLOC-based ELISA device can be used to evaluate drug toxicity in organoids.

## Discussion

The development of a practical MPS is a desirable alternative to animal experiments. In particular, MPSs that mimic multiple organs on a device are being developed to mimic the human body and to verify the response of each organ to drugs and the effects of interactions between organs. Kamei et al. have developed a microfluidic device with multiple valves in the flow paths connecting two culture chambers.^10, 26^ By controlling the valves, one can perform independent cell culture and immunostaining for each chamber or construct a closed circulation loop and perfuse the culture medium to evaluate the cardiotoxicity caused by drug metabolites in hepatocytes and analyze the interaction of the intestinal-liver axis. Shinha et al. have also developed a simple microfluidic plate (KIM-Plate) in which two chambers are connected by microchannels equipped with stirrers that allow cell seeding and medium exchange, using the same method as conventional well plates.^27^ However, these devices are designed to allow interaction between two different organs, making it difficult to perform multiorgan interactions. To replicate the interactions among multiple organs, a device that can connect four artificial human tissues was reported.^28^ The device was fabricated by inserting independently cultured tissue chips into a chamber that mimicked a blood vessel. They successfully perfused and cultured four tissues using this device. Another device succeeded in perfusion culture of 4–10 tissue cell types limited to planar culture.^29^ In this device, Transwells seeded with each type of tissue cells were placed in chambers connected by flow paths. By controlling multiple pneumatically driven pumps, independent flow control was possible for each chamber. However, these devices can deliver fluids in only one direction and are not designed to perform live imaging. Most importantly, these traditional devices are manufactured with pre-designed features (e.g., number, type, and arrangement of culture chambers, connecting channels, and analytical capabilities) and cannot accommodate operations outside their design.

In contrast, the proposed BLOC is characterized by its high versatility and scalability, allowing the construction of various types of MPSs according to purpose simply by connecting the Culture, Control, and Analysis BLOCs (Fig. 1). As the Culture BLOC can be connected after culturing cells or organoids under appropriate conditions for each, there is no limit to the number or types of cells or organoids that can be connected. In addition to the 2D-culture BLOC with a flat bottom and the Organoid-culture BLOC with a U-shaped bottom, we also prepared a Gel-Bottom BLOC, which contains a protein hydrogel with controlled stiffness and microtopography, making it possible to culture the cells in an ECM-mimicking environment (Fig. 2b,c). All Culture BLOCs could be subjected to live imaging of the internal cells and organoids. Additionally, the Control BLOCs enabled fluid transfer in both two and three dimensions (Fig. 2e–j, Supplementary Movies 1–4).

The adsorption/absorption of hydrophobic small molecules to PDMS is one of the concerns related to BLOCs; however, we succeeded in suppressing this by using a modification method that utilizes radiation-induced chemical reactions (Fig. 3a). We used highly penetrating γ-rays that can modify not only the entire bulk of BLOCs, but also many BLOCs at once. Oxidation and crosslinking of the PDMS backbone is a possible mechanism of suppressing adsorption/absorption. The ionizing radiations change the hydrophobic methyl groups (Si–CH_3_) into hydrophilic silanol groups (Si–OH), converting PDMS into a glass-like Si-O*_x_*-rich (where *x* = 3 or 4) structure through crosslinking.^30^ The inhibitory effect on adsorption/absorption was not significantly affected by an irradiation dose in the range of 500 kGy to 2,000 kGy, although the transparency in the visible light region of 400–800 nm gradually decreased depending on the irradiation dose (Fig. 3b). This is believed to be due to the formation of double bonds such as C=C and C=O by γ-ray irradiation. Thus, an irradiation dose of 500 kGy was considered optimal for suppressing the adsorption/absorption of small hydrophobic molecules and ensuring as much transparency as possible. We also confirmed that the PDMS modified with 500 kGy retained its original high oxygen permeability (Fig. 3c). Notably, the reaction range of widely used plasma is limited to a few hundred nanometers from the surface, whereas γ-rays enable bulk modification on the order of centimeters to meters. Therefore, unlike plasma treatment, in which molecular exchange occurs between the modified surface layer and the unmodified layer underneath, the modification effect of γ-rays lasts for a long period of time.^30^

The Culture BLOCs were confirmed to be suitable for cell culture because HSC-1 cells adhered to the bottom surface of the chambers with many remaining viable cells (Supplementary Fig. 2). However, the behavior of cells or organoids on PDMS substrates does not necessarily replicate their behavior *in vivo.* The material properties, elasticity, and surface topography of the substrate influence cell adhesion, morphology, and physiological functions.^18–20^ In fact, by controlling the elastic modulus of the protein hydrogels produced using the radiation-induced crosslinking technique, it was possible to culture HSC-1 cells not only as a monolayer but also as spheroids with a tumor-like structure *in vivo* (Fig. 4c,d). This difference occurs because cells spread out in a monolayer on softer substrates, i.e. 50 kPa hydrogel, owing to weaker cell-cell and cell-substrate adhesion forces. In contrast, cell-cell adhesion forces become stronger on substrates, such as 200 kPa hydrogels, with moderate stiffness than cell-substrate adhesion forces, leading to self-condensation.^31^ Additionally, the C2C12 myotubes aligned themselves along the direction of line patterning on the surface of the hydrogel, similar to the alignment of skeletal muscle fibers *in vivo* (Supplementary Fig. 3b). This phenomenon is attributed to mechanotransduction, whereby cells respond to microtopographical features of the substrate.^32^ Thus, incorporating a culture substrate such as a radiation-crosslinked hydrogel within the chip allows for the culture of cells and organoids in a manner that induces morphology and physiological functions more closely resembling those in the body.

In this study, we succeeded in connecting up to six Organoid-culture BLOCs and perfusing the solution. Any number of BLOCs can be connected. However, when perfusion is performed, it is not possible to pump solution to all connected BLOCs because of the pressure loss. The pressure loss depends on the type of used BLOCs (channel shape), the direction of connection, the type of fluid, and the pumping speed etc. The user can build up his/her own pressure loss based on these values. Users should consider whether perfusion with their own BLOC devices is feasible based on these values.

In this study, we evaluated the acute toxicity of acrylamide to human organoids by mimicking the circulatory system in the body after oral intake using a perfusion device constructed with Organoid-culture BLOCs. We were able to obtain live images of organoids during perfusion culture and analyze the viability of organoids against acrylamide over time using this device (Fig. 6b–d, and Supplementary Movies 5,6). The percentage of dead cells in the liver and heart organoids increased after 6 h perfusion, whereas the response in the kidney organoids was slower, starting at 12 h (Fig. 6c). Oral intake of acrylamide can damage the liver, kidneys, and brain, as well as cause damage to the central nervous system in experimental animals.^33, 34^ However, there are few reports of acute poisoning in humans owing to the large amount of oral intake. In the few case reports available, only central nervous system symptoms, such as impaired consciousness and convulsions, were observed owing to the rapid absorption of acrylamide into the body.^35, 36^ We confirmed that acrylamide directly affects organoids of the human liver, heart, brain, and kidneys and induces cell death. Although it is not possible to accurately determine whether the acute effects of acrylamide on the human body have been predicted, the use of MPS constructed with BLOCs suggests the degree of impact of acrylamide on the organs of the human body.

Analytical devices that can measure parameters such as protein expression, nucleic acid expression, and membrane potential using microsamples have been developed.^37–39^ These devices can automatically control small amounts of reagent, thereby preventing sample loss and contamination. As a result, they are highly compatible with MPS, which use small amounts of culture medium to culture cells and organoids. However, the MPS reported to date cannot be integrated into analytical devices and lack versatility. Therefore, we have developed BLOCs that not only have a Culture BLOC but also have functions for “Control” and “Analysis” (Fig. 2b–k). By attaching these BLOCs to the Culture BLOC, it is possible to construct a biochemical analysis device that enables various evaluations of organoids and culture supernatants during or after culture. For example, we developed a BLOC that enables ELISA and built an ELISA device that can assess the toxicity of chemical substances on heart organoids by measuring the amount of GAPDH released due to cell damage (Fig. 7a,e,f). As the device can be assembled simply by attaching and detaching BLOCs, it minimizes the negative effects (i.e., pressure) on the organoids and culture supernatant, thereby reducing the likelihood of human error. In the future, we will develop fluid-controlled BLOCs, such as pumps and solution mixing. In addition, we will develop BLOCs that can conduct various biochemical analyses such as immunostaining, PCR, and membrane potential measurements. Combining these BLOCs will allow building an MPS of greater versatility than conventional devices.

In this study, we developed BLOCs that can be used to freely configure various MPSs. All BLOCs, which have one of the functions of “Culture,” “Control” or “Analysis,” are standardized to the same size and can be connected in 2D or 3D using joints. In addition, to accurately evaluate the biological effects of the chemicals, we suppressed the adsorption of hydrophobic low-molecular-weight compounds to BLOC using a modification technique that utilizes radiation-induced reactions. Culture BLOC can be used not only for 2D cell culture and organoid culture but also for cultures that mimic the *in vivo* microenvironment by encapsulating radiation-crosslinked protein hydrogels with controlled stiffness and microtopography. By connecting multiple Culture BLOCs, we constructed a perfusion device for six organoids that mimicked systemic circulation and performed toxicity tests. Furthermore, by combining the “Culture,” “Control,” and “Analysis” BLOCs, we constructed an ELISA device that could successfully detect proteins released from the organoids.

Conventional MPSs can only be used for predetermined purposes, while the proposed BLOCs, can be used to freely configure MPSs according to specific purposes. In the future, we plan to create BLOCs with new functions to accommodate a wider range of culture conditions and analyses. For example, by changing the bottom layer of the Culture BLOC to glass, it is possible to perform highly accurate fluorescence imaging of cells and organoids using confocal microscopy. In addition, by using Culture BLOC, which is endowed with the control function of mechanical signals such as respiration and peristalsis, in addition to the introduction of an ECM-mimicking gel, it will be possible to reproduce organ functions to a higher degree. In the future, by simply combining these BLOCs like LEGO blocks, it will be possible to build an MPS quickly and cost-effectively, which will greatly contribute to basic research and preclinical trials.

## Methods

### Preparation of BLOCs

BLOCs were prepared by stacking multiple parts fabricated using soft lithography techniques (Supplementary Fig. 4). The molds for each part were fabricated using KT-0823-BK (Okamura Tech, Chiba, Japan) resin and a 3D printer (Smapri Sonic 4 K LL, Hotty Polymer, Tokyo, Japan. A φ5.0 mm metal sphere (67-2529-91, AS ONE, Osaka, Japan) was installed in the mold used for the bottom of the Organoid-culture BLOC. The hemisphere of the metal sphere was inserted into the mold. During the subsequent PDMS addition phase, the PDMS hardens along the hemisphere that wasn’t insetted into the mold; resulting in a U-shaped bottom.

Sylgard 184 (Dow, Midland, MI, USA), a mixture of base polymer and crosslinker at a ratio of 10:1, was poured into the mold and cured at 80 °C for 1 h. The cured PDMS was then removed from the mold to obtain the layers for BLOC construction. The basic BLOC structure was obtained by applying Sylgard 184 to the surface of each layer, stacking them together, and heating them at 80 °C for 1 h to bond the parts. In the case of the Check-valve BLOC, which controls the pumping direction, a φ1.0 mm stainless steel bead (17975168, MonotaRO, Osaka, Japan) was sealed in the channel before joining each part.

After the basic structure of the BLOC was modified using the radiation-induced reactions described later, convex or concave joints were attached to the designated points. The convex joints were made of a biocompatible resin (TrinDy DT-SG-160, Yamahachi Dental, Aichi, Japan) and fabricated using a 3D printer. The concave joints were composed of silicone tubes with outer and inner diameters of 2.0 and 1.0 mm, respectively (6-586-02, AS ONE). An adhesive (Super-X HyperWide, Cemedine, Tokyo, Japan) was used to bond the joints. Similarly, a silicone tube was bonded to an air vent at the top of the Culture BLOC.

To prevent leakage between the layers and at the joints, Sylgard 184 was applied to the outer circumference of the BLOC and cured at 80 °C for 1 h. Finally, two pillars made of KT-0823-BK and fabricated using a 3D printer were inserted at specified locations. This prevents the distortion of the BLOC shape when the chip is mounted on the frame described below.

The cap was fabricated using the same manufacturing and curing conditions as that followed for the BLOC parts. A silicon tube was attached to the cap of the BLOC tank. The pin that blocks the air vent was made with the 3D printer using KT-0823-BK as the resin. The top and bottom frames for fixing the BLOCs in position were fabricated using the 3D printer and STANDARD PHOTOPOLYMER RESIN Transparent Ib (FROGOODS, Kochi, Japan) as the resin. This resin is highly transparent in the visible light region and does not interfere with microscopic observations. PDMS sheet made of Sylgard 184 at a 100 μm thickness was adhered to the ground surface of the bottom frame. This step ensures increased friction of the bottom frame against the ground surface and prevents the movement of the BLOC-based device during operation. The PDMS sheet was fabricated in the same manner as the BLOC and caps. The bottom frame and PDMS sheet were bonded to the adhesive.

### Modification of PDMS utilizing radiation-induced chemical reactions

Using ionizing radiation-induced chemical reactions, we modified a BLOC made of PDMS. The modification effect was evaluated using 1 mm-thick Sylgard 184 sheets cured at 80 °C for 1 h. Sylgard 184 sheets in sealed bags were irradiated by ^60^Co γ-rays supplied from ^60^Co No.2 Irradiation Facility of Takasaki Institute in the air at ∼25 °C at a dose rate of 8 kGy/h (kGy = J/g). The adsorption/absorption behavior of the small hydrophobic molecules, optical transparency, and oxygen permeability of the samples before and after the γ-ray irradiation were evaluated.

The adsorption/absorption of small hydrophobic molecules in the samples was evaluated using fluorescent rhodamine B as a model drug.^16^ The sheet was fixed on the bottom of 35-mm tissue culture-treated polystyrene dishes (3000-35, AGC TECHNO GLASS, Shizuoka, Japan) using Kapton tape (650S-P, Teraoka Seiko, Tokyo Japan). After adding 4 mL of rhodamine B solution (250 ng/mL in deionized water) to the dishes, they were incubated at room temperature (∼25 °C) in the dark for 1 h. After incubation, the samples were washed thrice with 4 mL of deionized water and observed in 4 mL of deionized water. The absorption and distribution of rhodamine B in the samples were evaluated via confocal fluorescence imaging. Imaging was conducted using an upright microscope (BX51WI) with a disk scanning unit (BX-DSU), objective lens (LUMPLFLN40XW) (all from Evident, Tokyo, Japan), and CMOS camera (ORCA-Flash4.0 V3, Hamamatsu Photonics, Shizuoka, Japan). Image processing and analysis of fluorescence images were performed using ImageJ.^40–42^

A spectrophotometer (U3310, Hitachi, Ibaraki, Japan) was used to evaluate the transparency of the samples in the UV-visible light region. The samples were placed in a solid holder and measured at 25 °C at a wavelength range from 400 to 800 nm.

The oxygen permeabilities of the samples, sized Φ10 mm × 1 mm, were measured using a custom-built apparatus consisting of a glass vessel and a dissolved oxygen meter equipped with a diaphragm-type galvanic cell oxygen sensor (B-506S, Iijima Electronics Industry, Aichi, Japan). Briefly, the vessel was filled with Milli-Q water and saturated with nitrogen gas at a flow rate of 50 mL/min for 15 min to remove the dissolved oxygen. The oxygen concentration in the water was confirmed to be below the detection limit (0.01 mg/L) using a dissolved oxygen meter. To ensure uniform diffusion of dissolved oxygen, a stirrer was placed inside the vessel and stirred at 400 rpm. The concentration of oxygen that permeated the samples as a function of time was measured via contact with oxygen gas supplied from outside the vessel at 50 mL/min for 40 min. The diffusion coefficients of the sheets were estimated using Equation (1).

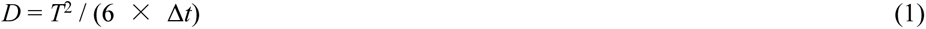

where *D* is the diffusion coefficient of oxygen gas, *T* is the thickness of the sheets, and Δ*t* is the delay time until oxygen permeated from the sheet is detected by the dissolved oxygen meter.

### Fabrication of ECM-mimicking hydrogels in Gel-bottom BLOCs

Acid-treated gelatin from porcine skin (639-37745, Nitta Gelatin, Osaka, Japan) was dissolved in deionized water (supplied by a Millipore Milli-Q system) at 10 or 15 wt% by heating at 50 °C for 30 min. After pouring the gelatin solution into the Culture BLOCs, the samples were sealed in plastic bags with an oxygen absorber (A-500HS, AS ONE) and stored overnight at 20 °C to undergo physical gelation. To produce micropatterned gelatin gels, PDMS molds were placed on top of the solution prior to physical gelation.

The PDMS molds were prepared as follows. The PDMS precursor (SIM-260) was mixed with a curing agent (CAT-260) (Shin-Etsu Chemical, Tokyo, Japan) in a 10:1 ratio. The mixed solution was evenly applied to a silicon master mold (DTM-1-1, Kyodo International, Kanagawa, Japan) and degassed. After curing at 150 °C for 30 min, the PDMS molds were peeled from the master mold.

The gelatin physical gels were irradiated using ^60^Co γ-rays at 15–20 °C at a dose rate of 4 or 5 kGy/h (kGy = J/g). This irradiation process transforms the physical gelatin gel into sterilized and crosslinked gelatin gels without any chemical treatment.^24^ After the PDMS molds were detached from the samples, the crosslinked gelatin hydrogels formed in Culture BLOCs were incubated with 150 μL of PBS overnight at 37 °C to remove non-crosslinked components and allow them to attain equilibrium. The Gel-bottom BLOC was used for cell culture after immersion in culture medium at 37 °C for 1 h to replace the absorbed PBS.

The compressive modulus of the hydrogels was measured using hydrogels formed on the bottom of 35-mm polystyrene dishes (3000-35, AGC Technoglass, Shizuoka, Japan) using the same method as described above. The measurements were performed at room temperature (∼25 °C) using a rheometer (RE2-33005C, Yamaden, Tokyo, Japan). The hydrogels were compressed with a 0.2 N load cell using a φ3 mm plunger at 50 µm/s, and the stress-strain curves were recorded. The obtained curves were then linearly fitted in the depth range less than 200 µm from the hydrogel surface to determine the compressive modulus.

### Construction and verification of various BLOC devices

Six Organoid-culture BLOCs with caps and vent closure pins were connected in a 2 × 3 configuration with joints and placed on the bottom frame to assemble a 2D perfusion device. A 50 mL syringe (1- 4908-08, Terumo, Tokyo, Japan) was filled with Milli-Q water colored with 2.0 mg/mL Food Red (4901325001245, Kyoritsu Foods, Tokyo, Japan) and connected to the inlet of the 2D perfusion device via a connecting silicone tube. The connecting silicone tube was made by bonding a silicone tube with outer and inner diameters of 2.0 and 1.0 mm, respectively, to a silicone tube with outer and inner diameters of 4.0 and 2.0 mm, respectively (1-596-05, AS ONE) with adhesive (Super-X Hyper Wide, Cemedine, Tokyo, Japan). The syringe was placed to a syringe pump (YSP-201, YMC, Kyoto, Japan), and the colored water was pumped into the perfusion device at 100 µL/min.

After all the Organoid-culture BLOCs were filled with colored water, one BLOC was removed and connected to three new Organoid-culture BLOCs with joints to form a 1 × 4 configuration. An empty 50 mL syringe was connected to the inlet of the device via a connecting silicone tube, and positive pressure was applied at 100 µL/min with a syringe pump.

A 3D perfusion device was constructed using two Organoid-culture BLOCs and two 3D-connector BLOCs in the upper and lower layers. The lower and upper layers were connected via a 3D-connect frame, which was bonded to the upper and lower frames with an adhesive in advance. Similar to the method followed for the 2D perfusion device, colored water was pumped into the upper or lower Organoid-culture BLOC at 100 μL/min.

A device for confirming the operation of the liquid flow direction control was constructed by connecting three Check-valve BLOCs and one Confluence BLOC. Three 50 mL syringes containing 200 μL of Milli-Q water colored with 2.0 mg/mL Food Red, Yellow, and Green (4901325001245, 4901325001146, 4901325000484, Kyoritsu Foods) were connected to each Check-valve BLOC. The three colored waters were pumped into the device in sequence at 100 µL/min using individual syringe pumps.

### Culture of cell lines in the Culture BLOCs

Human skin squamous cell carcinoma cell line (HSC-1; JCRB1015, JCRB Cell Bank, Osaka, Japan) was seeded into the 2D-culture BLOC with plasma treatment (YHS-R, SAKIGAKE-Semiconductor, Kyoto, Japan), and cultured in HSC-1 growth medium (Low-glucose DMEM, 11885-084, Thermo Fisher Scientific, Waltham, MA, USA) containing 20% FBS (12483020, Thermo Fisher Scientific), 2 mM L-Glutamine (G7513, Merck, Darmstadt, Germany), and 100 units/mL penicillin, 100 µg/mL streptomycin (15140122, Thermo Fisher Scientific) at 37 °C and 5% CO_2_ for 2 days. For live/dead imaging, the medium was replaced with HSC-1 growth medium containing 1 µg/mL Hoechst 33342 (H342, Dojindo Laboratories, Kumamoto, Japan), 1 µM calcein-AM (C396, Dojindo Laboratories), and 1 µg/mL PI (P378, Dojindo Laboratories) and incubated for 30 min. Phase contrast and fluorescence images of the cultured cells on the chip were acquired using an inverted microscope (IX83, Evident).

HSC-1 cells were seeded on the Gel-bottom BLOC in the same way as the 2D-culture BLOC and cultured in HSC-1 growth medium at 37 °C in 5% CO_2_ for 7 days. C2C12 myoblast cell line (EC91031101, DS Pharma Biomedical, Osaka, Japan) was seeded on the Gel-bottom BLOC and cultured in C2C12 growth medium (High-glucose DMEM, 08488-55, Nacalai Tesque, Kyoto, Japan) containing 10% FBS, 2 mM L-Glutamine, and 100 units/mL penicillin, 100 µg/mL streptomycin at 37 °C and 5% CO_2_. After seeding, myoblasts were switched to differentiation medium (High-glucose DMEM) containing 2% calf serum (16010167, Thermo Fisher Scientific), 2 mM L-Glutamine, 1× MEM Non-essential Amino Acids Solution (NEAA) (139-15651, FUJIFILM Wako Chemicals, Osaka, Japan), and 100 units/mL penicillin, 100 µg/mL streptomycin and cultured for 7 days. Phase-contrast images were obtained using a microscope (BZ-X800, Keyence, Osaka, Japan).

### Generation of hiPSC-derived organoids for toxicity assay

The protocol for generating fundic gastric organoid was based on established protocols.^43, 44^ Two days before the differentiation (day -2), hiPSCs (1383D2; HPS1005, RIKEN BRC, Ibaraki, Japan) in 0.5 mL of StemFit AK02N (Ajinomoto Healthy Supply, Tokyo, Japan) containing 10 µM Y27632 (036- 24023, FUJIFILM Wako Chemicals) and 1 µg of iMatrix-511 (892012, Matrixome, Osaka, Japan) were seeded into 24-well plate (3820-024, AGC Techno Glass) at a density of 10,000 cells per well. The next day (day -1), the medium was replaced with StemFit AK02N without Y27632. From day 0, the cells were differentiated in 0.5 mL of RPMI 1640 with 25 mM HEPES (22400089, Thermo Fisher Scientific), 100 units/mL penicillin, 100 µg/mL streptomycin and 100 ng/mL activin A (014-23961, FUJIFILM Wako Chemicals) for 3 days. Furthermore, 50 ng/mL BMP4 (020-18851, FUJIFILM Wako Chemicals) was added on day 0. FBS was added at 0.2% and 2% concentrations on days 1 and 2, respectively. From days 3 to 5, the medium was changed to RPMI 1640 with 25 mM HEPES, 2% FBS, 100 units/mL penicillin, 100 µg/mL streptomycin, containing 500 ng/mL FGF4 (065-06031, FUJIFILM Wako Chemicals), 200 ng/mL noggin (149-08861, FUJIFILM Wako Chemicals), and 2 µM CHIR99021 (034-23103, FUJIFILM Wako Chemicals) every day. On day 5, 2 µM retinoic acid (R2625, Merck) was added. On day 6, cells were dissociated using TrypLE (A1285901, Thermo Fisher Scientific) and seeded at 10,000 cells/well in V-bottom ultra-low attachment 96-well plates (PrimeSurface 96V Plate, Sumitomo Bakelite, Tokyo, Japan) in 100 µL of basal growth medium consisting of advanced DMEM-F12 medium (12634010, Thermo Fisher Scientific) with 1× B27 (w/o vitamin A, 12587010, Thermo Fisher Scientific), 1× N2 (17502048, Thermo Fisher Scientific), 2 mM GlutaMAX (35050061, Thermo Fisher Scientific), 100 units/mL penicillin, 100 µg/mL streptomycin, and 15 mM HEPES (15630080, Thermo Fisher Scientific), containing 100 ng/mL EGF (059-07873, FUJIFILM Wako Chemicals), 200 ng/mL noggin, 2 µM CHIR99021, 2 µM retinoic acid, and 10 µM Y27632. On day 9, the medium was changed to basal growth medium containing 100 ng/mL EGF, 200 ng/mL noggin, and 2 µM CHIR99021. On day 13, the medium was changed to basal growth medium containing 100 ng/mL EGF and 2 µM CHIR99021. The medium was changed every 3–4 days. On day 20, the medium was changed to basal growth medium containing 100 ng/mL EGF, 50 ng/mL FGF10 (069-06051, FUJIFILM Wako Chemicals), and 2 µM CHIR99021. The medium was changed every 2–3 days. On day 30, the medium was changed to basal growth medium containing 10 ng/mL EGF, 2 µM CHIR99021, 2 µM PD0325901 (162-25291, FUJIFILM Wako Chemicals), and 50 ng/mL BMP4. On day 32, the medium was changed to basal growth medium containing 10 ng/mL EGF, 50 ng/mL FGF10, and 2 µM CHIR99021. The medium was changed every 2–3 days. On day 36, the organoids were transferred to an ultra-low-attachment 24-well plate (3473, Corning, Corning, NY, USA). The medium was changed every 2–3 days.

The protocol for generating the intestinal organoids was based on previously established protocols.^45–47^ Two days before the differentiation (day -2), hiPSC in 0.5 mL of StemFit AK02N containing 10 µM Y27632 and 1 µg of iMatrix-511 were seeded into 24-well plate at a density of 10,000 cells per well. The next day (day -1), the medium was replaced with StemFit AK02N without Y27632. From day 0, the cells were differentiated in 0.5 mL of RPMI 1640 with 25 mM HEPES, 100 units/mL penicillin, 100 µg/mL streptomycin, and 100 ng/mL activin A for 3 days. FBS was added at 0.2% and 2% concentrations on days 1 and 2, respectively. From days 3 to 6, the medium was changed to RPMI 1640 with 25 mM HEPES, 2% FBS, 100 units/mL penicillin, 100 µg/mL streptomycin, containing 500 ng/mL FGF4 and 2 µM CHIR99021 every day. On day 7, cells were dissociated using TrypLE and seeded at 5,000 cells/well in V-bottom ultra-low attachment 96-well plates in 100 µL of basal growth medium consisting of advanced DMEM-F12 medium with 1× B27 (w/o vitamin A), 1× N2, 2 mM GlutaMAX, 100 units/mL penicillin, 100 µg/mL streptomycin, and 15 mM HEPES containing 100 ng/mL EGF, 500 ng/mL R-Spondin1 (181-02801, FUJIFILM Wako Chemicals), 100 ng/mL noggin, and 10 µM Y27632. On day 10, the medium was replaced with the basal growth medium containing 100 ng/mL EGF. The medium was changed every 2–4 days. On day 36, the organoids were transferred to an ultra-low-attachment 24-well plate. The medium was changed every 2–3 days.

The protocol for generating liver organoids was based on an established protocol for mixing hepatocytes, endothelial cells, and septum transversum mesenchyme (STM) cells.^48^ One day before differentiating the hiPSCs to hepatocytes, 600,000 hiPSCs in StemFit AK02N containing 10 µM Y27632 and 2.25 µg of iMatrix-511 were seeded into 35 mm culture dish (3000-035, AGC Techno Glass). Next day (day 0), the medium was changed to RPMI 1640 medium with 25 mM HEPES, 20% (v/v) StemFit For Differentiation (AS401, Ajinomoto Healthy Supply), 100 units/mL penicillin, 100 µg/mL streptomycin, and 100 ng/mL activin A. From day 0 to 3, 2 µM CHIR99021 was added to the medium. From day 1 to 4, 0.5 mM sodium butylate (193-01522, FUJIFILM Wako Chemicals) was added to the medium. The medium was changed every day. On day 6, the medium was changed to the medium consisting of StemFit AK02N without FGF2, 1% (v/v) dimethyl sulfoxide (D2650, Merck), 0.1 mM 2-mercaptoethanol (198-15781, FUJIFILM Wako Chemicals), 1 mM GlutaMAX, 1× NEAA, 100 units/mL penicillin, and 100 µg/mL streptomycin and cultured for 4 days. The medium was changed every day. One day before differentiating the hiPSCs to endothelial cells, 150,000 hiPSCs in StemFit AK02N containing 10 µM Y27632 and 2.25 µg of iMatrix-511 were seeded into a 35 mm culture dish. Next day (day 0), the medium was changed to DMEM-F12 medium with GlutaMAX (10565018, Thermo Fisher Scientific) with 20% (v/v) StemFit For Differentiation, 100 units/mL penicillin, 100 µg/mL streptomycin, 8 µM CHIR99021, and 25 ng/mL BMP4 and cultured for 3 days. The medium was changed every day. From day 3, cells were cultured in StemPro-34 SFM (10639011, Thermo Fisher Scientific) with 2 mM GlutaMAX, 100 units/mL penicillin, 100 µg/mL streptomycin, 200 ng/mL VEGF (226-01781, FUJIFILM Wako Chemicals), and 2 µM forskolin (F0855, Tokyo Chemical Industry, Tokyo, Japan) for 4 days. The medium was changed every day. On day 7, the cells were dissociated using 0.05% trypsin (Merck, T3924) and seeded on four 60 mm dishes (3010-060, AGC Techno Glass) precoated with 1 µg/cm^2^ fibronectin (FC010, Merck) and cultured for 2 days in StemPro-34 SFM with 2 mM GlutaMAX, 100 units/mL penicillin, 100 µg/mL streptomycin, and 50 ng/mL VEGF. Four days before differentiating the hiPSCs to STM (day -4), 29,000 hiPSCs in StemFit AK02N containing 10 µM Y27632 and 2.25 µg of iMatrix-511 were seeded into a 35 mm culture dish. The next day (day -3), the medium was replaced with StemFit AK02N without Y27632. From day 0, the cells were cultured in DMEM-F12 medium with GlutaMAX with 20% (v/v) StemFit for Differentiation, 100 units/mL penicillin, 100 µg/mL streptomycin, 8 µM CHIR99021, and 25 ng/mL BMP4 for 4 days. From day 1, 2 ng/mL activin A and 10 ng/mL PDGF-BB (160-19741, FUJIFILM Wako Chemicals) were added to the medium. The medium was changed every day. From day 4, the cells were cultured in StemPro-34 SFM with 2 mM GlutaMAX, 100 units/mL penicillin, 10 ng/mL FGF2 (064-04541, FUJIFILM Wako Chemicals), and 10 ng/mL PDGF-BB for 3 days. The medium was changed every day. Hepatocytes and STM cells were dissociated using 0.5× TrypLE and endothelial cells were dissociated using 1× TrypLE, and 30,000 hepatocytes, 21,000 endothelial cells, 6,000 STM cells were seeded in V-bottom ultra-low attachment 96-well plates in 150 µL of the medium consisting of 50% (v/v) High-glucose DMEM and 50% (v/v) KBM-VEC1 basal medium (16030110, Kohjin Bio, Saitama, Japan) supplemented with 2.5% (v/v) StemFit For Differentiation, 1 mM GlutaMAX, 100 units/mL penicillin, 100 µg/mL streptomycin, 0.1 µM dexamethasone (041- 18861, FUJIFILM Wako Chemicals), 10 ng/mL oncostatin M (152-03411, FUJIFILM Wako Chemicals), and 10 µM Y27632. Next day, 50 µL of the medium was removed and 100 µL of the medium without Y27632 was added. Half of the medium was replaced daily.

Generating heart organoids was based on an established protocol.^49^ Two days before differentiation (day -2), hiPSCs were seeded at 10,000 cells/well in V-bottom ultra-low-attachment 96-well plates in 100 µL of StemFit AK02N containing 10 µM Y27632. Next day (day -1), 50 µL of the medium was removed and 200 µL of StemFit AK02N without Y27632 was added. On day 0, 166 µL of the medium was removed and 166 µL of RPMI 1640 medium (11875093, Thermo Fisher Scientific) with 1× B27 (w/o insulin, A1895601, Thermo Fisher Scientific), 100 units/mL penicillin, 100 µg/mL streptomycin (RPMI/B27(-)/PS) containing 6 µM CHIR99021, 1.875 ng/mL BMP4 (314- BP-010, R&D Systems, Minneapolis, MN, USA) and 1.5 ng/mL activin A was added. After 24 h, 166 µL of the medium was removed and 166 µL of RPMI/B27(-)/PS was added. After 24 h, 166 µL of the medium was removed and 166 µL of RPMI/B27(-)/PS containing 3 µM Wnt-C59 (S7037, Selleck Chemicals, Houstan, TX, USA) was added. After 48 h, 166 µL of the medium was removed and 166 µL of RPMI/B27(-)/PS was added. On day 6, 166 µL of the medium was removed and 166 µL of RPMI 1640 medium with 1× B27 (17504044, Thermo Fisher Scientific), 100 units/mL penicillin, 100 µg/mL streptomycin (RPMI/B27(+)/PS) was added. On day 7, 166 µL of the medium was removed and 166 µL of RPMI/B27(+)/PS with 3 µM CHIR99021 was added. After 1 h of incubation at 37 °C with 5% CO_2_, the organoids were transferred to ultra-low attachment 24-well plate with 0.8 mL of RPMI/B27(+)/PS. The medium was changed every 3–4 days.

The protocol for generating brain organoids was based on an established protocol.^50^ On day 0, hiPSCs were seeded at 9,000 cells/well in V-bottom ultra-low attachment 96-well plates in 100 µL of the differentiation medium consisting of Glasgow’s minimal essential medium (12965-65, Nacalai Tesque) supplemented with 1× NEAA, 20% (v/v) knockout serum replacement (10828028, Thermo Fisher Scientific), 1 mM sodium pyruvate (P8574, Merck), 0.1 mM 2-mercaptoethanol, 100 units/mL penicillin, and 100 mg/mL streptomycin, 5 µM SB431541 (192-16541, FUJIFILM Wako Chemicals), and 3 µM IWR-1 (I0161, Merck) with 20 µM Y27632. On day 3, 100 µL of the differentiation medium with 20 µM Y27632 was added. From day 6, half of the medium was replaced with differentiation medium without Y27632 every 3 days. On day 18, the organoids were transferred to ultra-low attachment 24-well plate with 0.8 mL of culture medium consisting of DMEM-F12 medium with GlutaMAX, 1× N2, 1% chemically defined lipid concentrate (11905031, Thermo Fisher Scientific), 100 units/mL penicillin, 100 µg/mL streptomycin, and 250 ng/mL amphotericin B (15290018, Thermo Fisher Scientific). The medium was changed every 3–4 days. From day 35, 10% (v/v) FBS and 5 µg/mL heparin (H4784, Merck) were added to the culture medium.

The protocol for generating kidney organoids was based on the established protocols.^51–53^ One day before the differentiation (day -1), 135,000 hiPSCs in StemFit AK02N containing 10 µM Y27632 and 2.25 µg of iMatrix-511 were seeded into a 35 mm culture dish. On day 0, the medium was changed to STEMdiff APEL2 medium (05270, STEMCELL Technologies, Vancouver, Canada) with 5% PFHM-II protein-free hybridoma medium (12040077, Thermo Fisher Scientific), 100 units/mL penicillin, 100 µg/mL streptomycin (APEL2/PFHM-II/PS) containing 8 µM CHIR99021. The medium was changed every 2 days. On day 4, the medium was changed to APEL2/PFHM-II/PS containing 1 µM CHIR99021, 200 ng/mL FGF9 (066-06201, FUJIFILM Wako Chemicals), and 1 µg/mL heparin. The medium was changed every 2 days. On day 7, the cells were dissociated using TrypLE and seeded at 25,000 cells/well in V-bottom ultra-low attachment 96-well plates in 100 µL of Essential 6 medium (A1516401, Thermo Fisher Scientific) with 100 units/mL penicillin, 100 µg/mL streptomycin (E6/PS) containing 1 µM CHIR99021, 200 ng/mL FGF9, and 1 µg/mL heparin. Next day, 100 µL of the medium was added. From days 9–11, half of the medium was replaced daily. On day 12, the organoids were transferred to an ultra-low-attachment 24-well plate filled with 0.8 mL E6/PS. The medium was changed every 3 days.

### Functional evaluation of perfusion device

Six Organoid-culture BLOCs and their caps were sterilized by using γ-ray irradiation at the dose of 15 kGy. The bottom frame, top frame, and air vent-closing pins were placed in an ozone sterilizer (CoolCLAVE Blue Ozone and UV Sterilizer, FirstResponder Technologies, San Diego, CA, USA) and irradiated with ozone/UV light for 30 min. A perfusion device was constructed by connecting the Organoid-culture BLOCs through joints to form a 2 × 3 configuration. The constructed device was placed in the bottom frame. Three organoids from each organ prepared as described above were added to each BLOC. Organoid species were applied from the inlet to the outlet: stomach, intestines, liver, heart, brain, and kidneys.

Three of each organoid species were transferred to each Organoid-culture BLOC filled with perfusion medium consisting of advanced DMEM-F12 medium with 1× B27 (w/o vitamin A), 1× N2, 2 mM GlutaMAX, 100 units/mL penicillin, 100 µg/mL streptomycin, and 15 mM HEPES. Perfusion medium was added so that the amount of medium in the chamber of the culture BLOC was approximately 150 µL. The chamber of the Culture BLOC was then closed with a cap and air vent closure pin. The Perfusion culture devices were mounted on an inverted microscope. The temperature and CO_2_ were set to 37 °C and 5%, respectively, adjusted using a stage-top incubator (STXF-IX3WX, Tokai Hit, Shizuoka, Japan). For live/dead imaging, a 50 mL syringe filled with perfusion medium with 1 µg/mL Hoechst 33342 and 1 µg/mL PI was connected to the device inlet through a connecting silicone tube. The connected syringe was placed in a syringe pump and the perfusion culture was performed by pumping fluid into the device at the flow rate of 10 μL/min for 24 h. For toxicity assay, 30 mM acrylamide (019-08011, FUJIFILM Wako Chemicals) was added to the perfusion medium in the syringe. Time-lapse images were captured hourly using an objective lens (UPLFLN4XPH, Evident) and a CMOS camera (Zyla4.2 PLUS, Andor, Belfast, Northern Ireland).

### Construction and utilization of the ELISA device

Human iPSC-derived heart organoids were placed in Organoid-culture BLOCs filled with 150 µL advanced DMEM-F12 with 1× B27 (w/o vitamin A) containing 1× N2, 2 mM GlutaMAX, 100 units/mL penicillin, 100 µg/mL streptomycin, and 15 mM HEPES and cultured for 24 h in an incubator at 37 °C with 5% CO_2_. For the toxicity assay, 30 mM of acrylamide was added. After incubation, the Culture BLOC was sealed with a cap and closure pin.

Before ELISA device construction, the Tank BLOCs were filled with 250 μL of 1% BSA in PBS or 100 μL of Alexa Fluor 488-conjugated GAPDH secondary antibody (1:100, #3906, Cell Signaling Technology) in 1% BSA in PBS and then sealed with a cap and closure pin. Thereafter, 40 µL of PBS containing 1 µL of Total GAPDH MAPmate Magnetic Beads (46-667MAG, Merck) was applied to the ELISA BLOC after attaching a neodymium magnet (NOS093, NeoMag, Tokyo, Japan) to the bottom. The ELISA BLOC was left to stand at room temperature for 1 h.

The ELISA device was constructed using the Tank BLOCs, Organoid-culture BLOC, Check-valve BLOC, Confluence BLOC, and ELISA BLOC, which were operated as described above (Fig. 7a). The BLOCs were connected by a joint, and the Tank BLOC and the Organoid-culture BLOC were each connected to ring pumps (RP-QⅢB1.5S-P35C-DC3VS, Aquatech) that were connected to a controller unit (RE-C500, Aquatech, Osaka, Japan). The liquid delivery control program for the controller unit was written on a laptop (PC-GN186RULH, NEC, Tokyo, Japan).

ELISA was performed using the constructed device. The ring pump A was operated at 100 µL/min for 2 min to pump culture supernatant from the Culture BLOC to the ELISA BLOC, and then allowed to stand for 2 h at room temperature (∼25 °C). Then, ring pump B was operated at 100 µL/min for 5 min to pump 1% BSA in PBS from the Tank BLOC to the ELISA BLOC. Next, ring pump C was operated at 100 µL/min for 1.5 min to pump the secondary antibody solution from the Tank BLOC to the ELISA BLOC, and then allowed to stand for 1 h at room temperature. Finally, ring pump B was driven at 100 µL/min for 5 min to pump 1% BSA in PBS from the Tank BLOC to the ELISA BLOC.

Fluorescence images of the magnetic beads were captured using an inverted microscope with an objective lens (UPLFLN10XPH2, Evident) and an EMCCD camera (iXon Life 888, Andor). A U-FBNA mirror unit (Evident) was used to observe Alexa Fluor 488. For the red-fluorescent magnetic beads, an excitation filter (FF01-624/40, Semrock), dichroic mirror (FF660-Di02, Semrock), and emission filter (FF01-692/40, Semrock) were used. The green and red fluorescence intensities of each bead were measured using the ImageJ software. The green/red ratio was calculated to compensate for the variation in intensity due to differences in focus.^54^

### Statistics

Error bars represent the mean ± standard error of the mean (SEM). An unpaired two-sided Student’s *t*-test was performed to evaluate statistical differences between the two groups, and values of *P* < 0.05 were considered statistically significant. The analysis was performed in Excel for Microsoft 365 MSO (v2049, Microsoft, Redmond, WA, USA). Values of *P* and *N* used to calculate the statistics were described in the figure legends.

## Acknowledgments

The authors thank Ms. Ryoko Mezaki, Ms. Yukiko Sakatsume, Ms. Noriko Tawara, Ms. Emika Taguchi, Ms. Naoko Kono, and Dr. Zorigt Odgerel for technical assistance. The authors thank Dr. Yuji Morimoto for providing the HSC-1 cell line. We would like to thank Editage for the English language editing. This study was supported by JSPS KAKENHI (Grant Nos. 22H05054, 23K17190 and 24K01998) and ACT-X (Grant No. JPMJAX2014), and A-STEP (Grant No. JPMJTR22U7) from the Japan Science and Technology Agency (JST), Innovative Science and Technology Initiative for Security of Acquisition, Technology, and Logistics Agency (Grant No. JPJ004596).

## Author contributions

Y.K., H.H., K.O., T.G.O., N.I., and M.T. designed the study. Y.K., H.H., K.O., T.G.O., A.K., and Y.O. performed the experiments and analyzed the data. All the authors have written and approved the final manuscript.

