## Supplementary Information for "BLOC: buildable and linkable organ on a chip"

**Supplementary Figures**

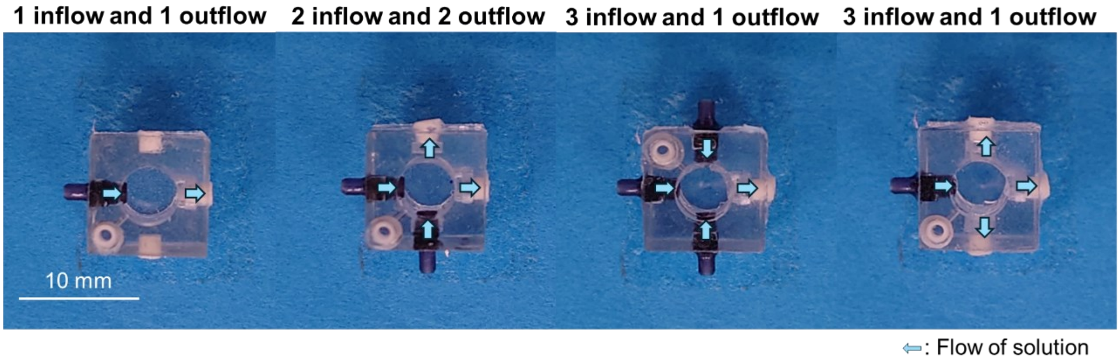


**Supplementary Figure 1. Variation of flow patterns in Culture Buildable and Linkable Organs on a Chip (BLOCs).** Pictures show the top views of the Culture BLOCs.

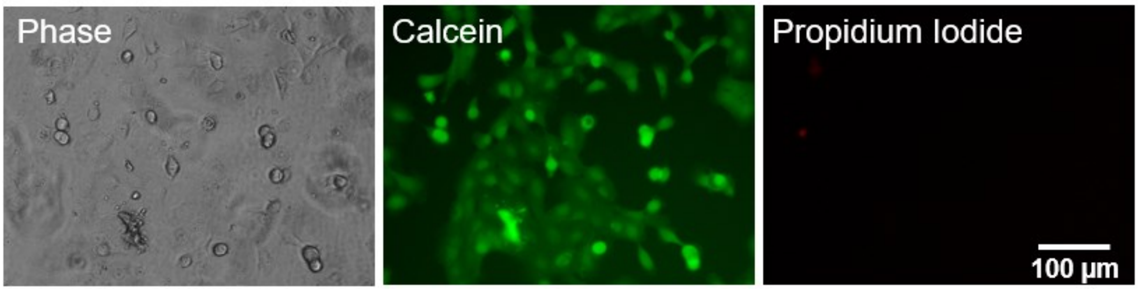


**Supplementary Figure 2. HSC-1 cells cultured in the 2D-culture BLOC for 48 h.** Left, phase contrast image. Center and right fluorescence images show the cells stained by calcein (green) and propidium iodide (red), respectively.

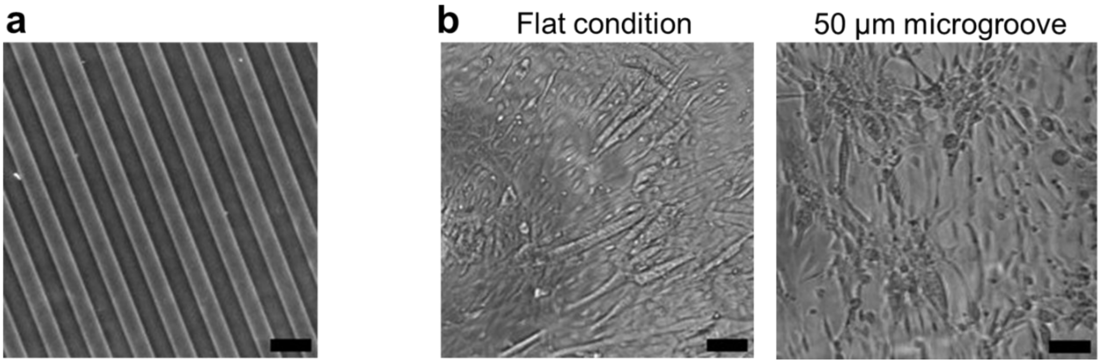


**Supplementary Figure 3. Microfabricated hydrogel in the Gel-bottom BLOC.** (a) Phase contrast image of gelatin hydrogel fabricated with 50 μm microgroove on the surface. (b) Phase contrast images of C2C12 myoblasts cultured on the hydrogel in Gel-bottom BLOCs for 5 days under flat condition (left) or 50 μm microgroove (right). Scale bars, 100 μm.

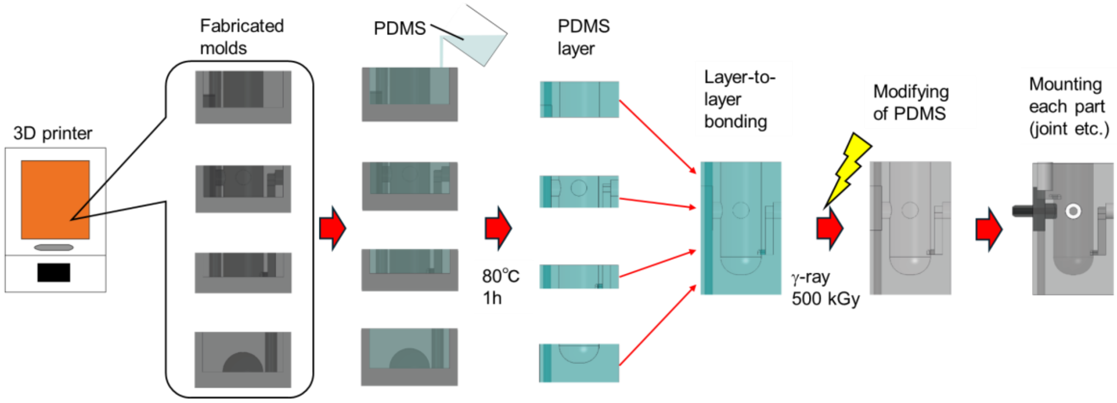


**Supplementary Figure 4. Fabrication method of BLOCs.** PDMS was poured into a mold fabricated by the 3D printer and heated at 80 °C for 1 h to cure the PDMS. After curing, the fabricated PDMS layers were removed from the mold and bonded to each other by applying PDMS to the surface of each layer part and heating them at 80 °C for 1 h. The bounded parts were modified by irradiating them with 500 kGy of γ-rays. Necessary parts for the BLOC, such as convex joint and concave joints, were attached to the modified parts. PDMS was then applied to the outer circumference of the BLOC and cured at 80 °C for 1 h and the BLOC was completed.

**Supplementary Movies**

**Supplementary Movie 1. 2D pumping experiment with the constructed perfusion device.** This movie was taken at 30 fps and speeds of up to 10×.

**Supplementary Movie 2. 2D pumping experiments with the perfusion device reconstructed after the pumping experiments.** This movie was taken at 30 fps and speeds of up to 10×.

**Supplementary Movie 3. 3D pumping experiment with the constructed perfusion device.** This movie was taken at 30 fps and speeds of up to 10×.

**Supplementary Movie 4. Pumping experiment with the constructed check-valve device.** This movie was taken at 30 fps and speeds of up to 15×.

**Supplementary Movie 5. Co-culture of human organoids connected in the perfusion device.** Time-lapse fluorescence imaging of six kinds of organoids derived from hiPSCs in the presence of Hoechst stain (cyan) and propidium iodide (red) for 24 h without toxic agents. The perfusion flow originated from the stomach (top left), intestine (top middle), liver (top right), heart (bottom right), brain (bottom middle), and kidney (bottom left). Scale bar, 1 mm.

**Supplementary Movie 6. Toxicity assay of human organoids connected in the perfusion device.** Time-lapse fluorescence imaging of six kinds of organoids derived from hiPSCs in the presence of Hoechst stain (cyan), propidium iodide (red), and 30 mM acrylamide for 24 h. The perfusion flow originated from the stomach (top left), intestine (top middle), liver (top right), heart (bottom right), brain (bottom middle), and kidney (bottom left). Scale bar, 1 mm.
